# Megaherbivore mortality risk from temperature extremes is buffered by vegetation productivity but amplified by rainfall

**DOI:** 10.64898/2026.05.29.728837

**Authors:** Hansraj Gautam, Alexandre Courtiol, John Jackson, Mirkka Lahdenperä, Martin Seltmann, Axelle Delaunay, Diogo F. dos Santos, Zaw Min Oo, Win Htut, Virpi Lummaa

**Affiliations:** Department of Biology, University of Turku, Turku, Finland 20014; Department of Evolutionary Genetics, Leibniz Institute for Zoo and Wildlife Research, Alfred-Kowalke-Strasse 17,10315 Berlin, Germany; Estación Biológica de Doñana (EBD-CSIC), Seville, Spain, 41092; Myanma Timber Enterprise, Ministry of Natural Resources and Environmental Conservation, Yangon 11011, Myanmar

**Keywords:** Climate change, megafauna, demography, heat, vegetation productivity, ecological buffer

## Abstract

Demographic risks from climate change remain poorly understood for megaherbivore species which face unique challenges because their large food requirements impose strong bottom-up limitation while their small surface-area-to-volume ratio limits heat dissipation. Here, we test how megaherbivore mortality is shaped by the concurrent effects of temperature and rainfall along with vegetation productivity, a potentially critical modifier of climate effects. By analysing four decades of monthly mortality records from 4,457 semi-captive Asian elephants across Myanmar, we identify multiple environmental pathways regulating megaherbivore mortality. Elephant mortality increased at extreme hot and cold temperatures in regions with low annual vegetation productivity, whereas high vegetation productivity buffered against such U-shaped effects of temperature. Furthermore, high rainfall amplified the negative impacts of extreme heat, underlining risks arising from the joint effects of heat and humidity. Seasonal declines in vegetation productivity did not explain the elevated mortality at temperature extremes. Together, our findings show that megaherbivores face elevated risks from global warming, but such risks strongly depend on vegetation productivity and humidity, highlighting multiple pathways through which climate change can shape the dynamics of megaherbivore populations.

## INTRODUCTION

Effectively mitigating the risks posed by climate change to animal populations requires understanding their demographic responses to environmental variation (Thomas et al. 2004; Orgeret et al. 2022; Spooner et al. 2018; Paniw et al. 2021). Such responses are reliably understood by analysing vital rates quantified from individual-based demographic records (Paniw et al. 2022; Thorley et al. 2025; Woodroffe et al. 2017), but this knowledge is increasingly poor along the body size continuum as it is difficult to monitor the multi-decade life-histories of large animals (Fuller et al. 2016). Addressing this empirical gap is crucial to manage threats to megafauna (weighing over 60 kg) which have oversized roles in ecosystem functioning (Pringle et al. 2023; Malhi et al. 2022) but their already vulnerable populations are facing heightened risks from climate change (Duncan et al. 2012; Ogutu and Owen-Smith 2003; Ripple et al. 2015; Hetem et al. 2014).

The demographic impacts of climate change can manifest through multiple pathways (Fuller et al. 2021; Ickin et al. 2025; Paniw et al. 2021). First, rainfall can strongly determine population growth by shaping the availability of key resources impacting survival and fertility (Coe et al. 1976; Stommel et al. 2016; Western et al. 2015; Ogutu and Owen-Smith 2003). Second, changes in the ambient temperature can also impact survival and reproduction, although empirical studies on megafauna remain scarce (Fuller et al. 2016; Hetem et al. 2014; Walsh et al. 2019). Furthermore, perhaps due to the popular emphasis on global warming, studies on tropic-dwelling taxa often focus on impacts at the hot end of temperature regimes while the cold end is rarely examined (Spooner et al. 2018; Veldhuis et al. 2019). Changes at the cold end can nonetheless impact animal performance, especially in endotherms that need to regulate body temperatures within narrow ranges (Boyles et al. 2011; Cordes et al. 2020; Masoero et al. 2020; Vetter et al. 2015). Third, the interactive effects of temperature and rainfall can produce divergent demographic outcomes. For example, heat can amplify mortality risk induced by droughts (Fuller et al. 2021; Thorley et al. 2025), whereas high humidity can amplify heat stress by constraining evaporative heat loss (Raymond et al. 2020; Coulson et al. 2025). Therefore, considering concurrent effects of multiple stressors can better inform our knowledge of risks from climatic disruptions.

It is also challenging to disentangle whether the apparent impacts of climatic disruptions resulted from the effects of food scarcity (Creel et al. 2023; Fuller et al. 2021). For example, earlier studies attributed the decline in African wild dog populations to reduced survival and reproductive success under heat stress (Woodroffe et al. 2017; Rabaiotti et al. 2023), but a recent study argued that prey scarcity had greater impacts (Creel et al. 2023), inviting criticism for downplaying the threats from global warming (Woodroffe et al. 2023). This “hot vs. hungry” debate applies to several taxa, given the complex ways in which food shortage can shape the effects of temperature extremes (Vetter et al. 2015; Fuller et al. 2021). For instance, starvation can make thermoregulation inefficient, increasing mortality risk (Rey et al. 2017; Fuller et al. 2021), whereas demographic impacts of hot conditions may not be apparent if high vegetation productivity buffers heat stress by increasing food abundance (Thorley et al. 2025). This complexity underscores inferential limits of considering temperature effects in isolation, particularly in species with large food requirements as both low vegetation productivity and climate extremes can affect their demography (Mduma et al. 1999; Western et al. 2015; Ogutu and Owen-Smith 2003).

Megaherbivores, weighing over 1000kg, may be uniquely sensitive to climate because their large body size strongly shapes both trophic regulation and thermoregulatory constraints. First, due to their massive food requirements and the near absence of natural predation in adults, megaherbivore populations are said to be primarily bottom-up regulated by food availability (Owen-Smith 1988; Pringle et al. 2023). Their body condition, survival and distribution strongly depends on forage quantity and quality [e.g. black and white rhinos (Ferreira et al. 2019; Ndlovu et al. 2023), giraffe (Strauss et al. 2015), African forest elephants (Bush et al. 2020), savannah elephant (Trimble et al. 2009), multiple species (Abraham et al. 2025)]. Such bottom-up regulation can shape the climatic control of population dynamics, for example, through effects of starvation on thermoregulation (Fuller et al. 2016). Second, most megaherbivores may be vulnerable to temperature extremes because their relatively low surface-area/body-volume ratio limits dissipation of body heat (Weissenböck et al. 2012; Dunkin et al. 2013). As vegetation productivity can modify the impact of temperature extremes by influencing forage and shade availability, considering its effects on demography can shed light on climate-linked threats to megaherbivores.

This paper investigates the concurrent effects of temperature, rainfall and vegetation productivity in the Asian elephant, a megaherbivore with profound role in ecosystem functioning (Sukumar 2003; Campos-Arceiz and Blake 2011). While such long-lived species with a slow pace-of-life are said to be generally buffered against short-term climatic disruptions (Morris et al. 2008), a comprehensive understanding is still developing for the three extant elephant species. Studying elephants can offer crucial insights about demographic processes in distinct habitats of megaherbivores, because their distribution ranges show large inter-species differences in environmental stressors (Dunkin et al. 2013; Sukumar 2003). Most past studies have focused on the semi-arid and open canopy habitats where savannah elephants occur, whereas limited understanding exists for the hot, humid and dense-canopy niches occupied by the African forest elephant and the Asian elephant. Even in the case of savannah elephants, studies have often relied on coarse datasets obtained from surveys conducted by park management (Trimble et al. 2009; Wato et al. 2016; Duncan et al. 2012). However, as such long-lived species exhibit a large age-dependent variation in mortality and fertility (Lee et al. 2011; Lahdenperä et al. 2018), demographic consequences of climate are more reliably understood from individual-based records. For example, long-term monitoring of elephants in Amboseli, Kenya, has revealed not only immediate negative impacts of droughts on conceptions and survival, but that early-life experience of droughts can also compromise survival and reproduction in later ages (Lee et al. 2022; 2011; 2013), effects that appear to be linked with forage availability (Boult et al. 2018; Wato et al. 2016; Rasmussen et al. 2006). However, a critical missing piece in this knowledge is how these populations responded to temperature regimes.

To advance this understanding, we analysed four decades of mortality data derived from individual-based records of semi-captive Asian elephants in Myanmar, representing the world’s largest semi-captive elephant population occupying hot and humid habitats. Detailed records of these elephants maintained by the Myanma Timber Enterprise (MTE) offer rich long-term demographic data (Mar et al. 2012). This dataset is particularly suited to analyse mortality because ages are accurately known for most elephants, and the exceptionally large sample size it offers is essential to quantitatively model mortality in such species with overall high survival. Crucially for our study, the wide geographical spread of these elephants facilitates statistically powerful analyses of the environmental drivers of mortality across a wide bioclimatic niche (12 administrative divisions of Myanmar, henceforth called divisions, Fig 1, SI Fig S2). These attributes of demographic data are very difficult to achieve for wild populations. While these elephants are used in timber extraction activities for an average five to six hours a day, they spend most of the day foraging on naturally available plants in the same forests used by wild elephants (Mar et al. 2012; Dierenfeld et al. 2020). Therefore, the effects of environment and habitat conditions on them are expected to be closely comparable to wild elephants and distinct from captive elephants in zoos, as indicated by mortality rates that are closer to wild populations rather than zoo elephants (De Silva et al. 2013; Mar et al. 2012). Advancing the previous considerations of climatic predictors in isolation, we here examine vegetation productivity as a key predictor as its potential bottom-up effects can shed light on the survival consequences of temperature regimes seen in this population. Previously, Mumby et al. (2013) analysed a smaller dataset restricted to two divisions and found substantially higher mortality in both hot and cold conditions, relative to monthly average temperatures of ∼24 °C which supported highest survival. We hypothesized that high mortality in hot and cold months could be underpinned by seasonal drops in vegetation productivity. Furthermore, we expected that elephants in habitats with greater vegetation productivity would be buffered against stress from extreme temperatures, because of forage and shade advantage. We examined these two potential explanations by isolating the effects arising from the spatial differences across divisions in annual vegetation productivity as captured by remotely sensed Normalized Difference Vegetation Index, i.e., NDVI (Pettorelli et al. 2011) and its seasonal fluctuations (Methods). Specifically, we tested seven predictions:

**Figure 1.**
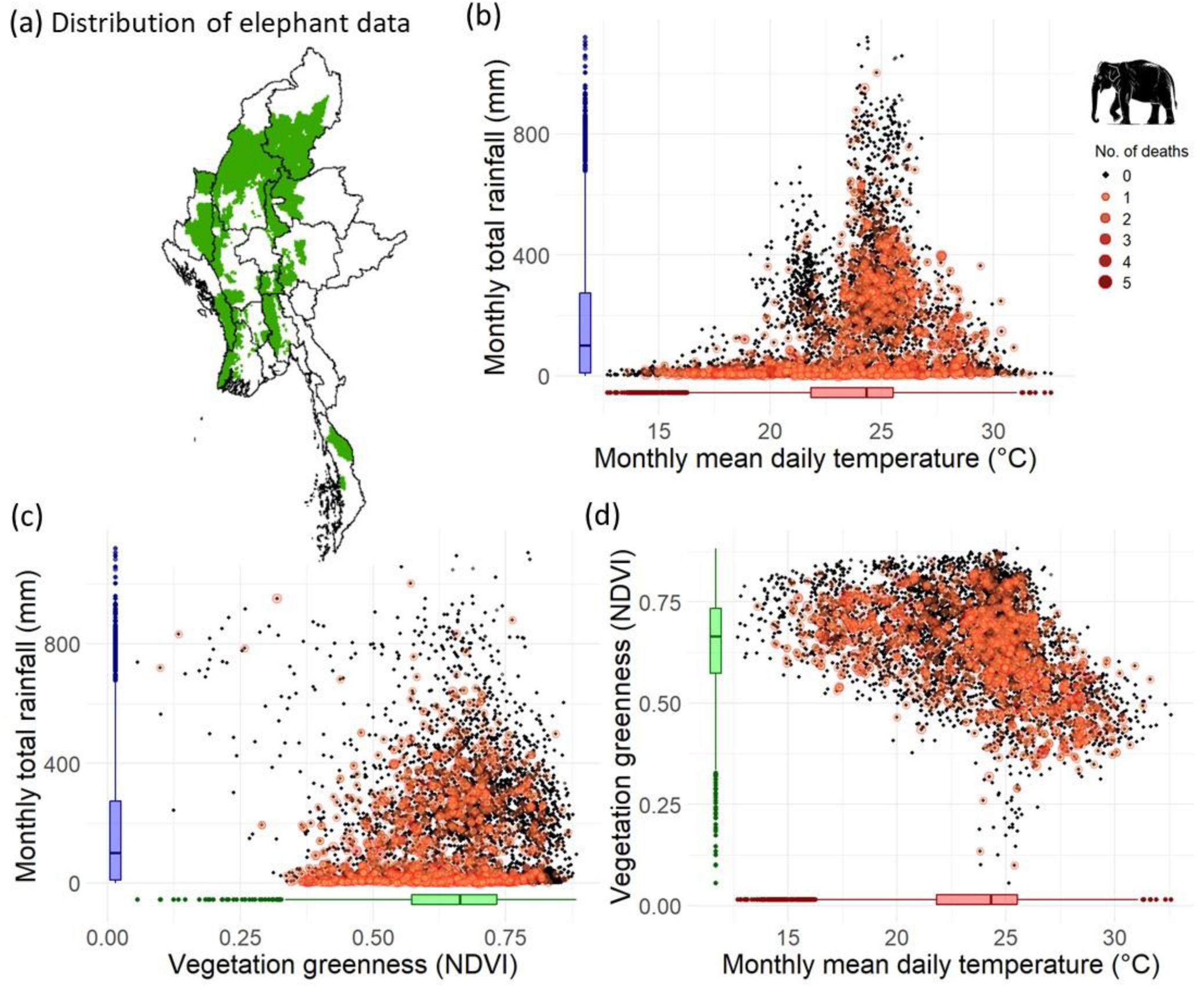
The wide environmental niche covered in this study a) Distribution of demographic data in different administrative divisions (see also Fig. S1), and distribution of elephant deaths along: b) temperature and rainfall, c) NDVI and rainfall, and d) temperature and NDVI. Data points represent monthly within-division values during 1982–2020 and are pooled across all age and sex classes analysed (Methods). Box-plots show the distribution of the environmental variable.

***P1***. Low rainfall increases mortality.

***P2***. Extreme temperatures increase mortality (i.e., U-shaped effects of temperature).

***P3***. Low rainfall intensifies effects of extreme temperatures (interaction effects).

***P4***. Divisions with higher vegetation productivity have lower mortality.

***P5***. Divisions with higher vegetation productivity are buffered against the effects of extreme temperatures (interaction effects).

***P6***. Seasonal declines in vegetation productivity increase mortality.

***P7***. The effects of extreme temperatures weaken after accounting for the effects of vegetation seasonality (both additive and interaction effects).

## RESULTS

### Demographic and environmental data

We used life records of 4,457 elephants to prepare a time series of monthly mortality, the binomial response variable which was specified as a two-column matrix of number of deaths and total elephants present, for each of the 12 administrative divisions, spanning January 1982 to December 2020 (Fig. 1, Methods). These time series were obtained for 10 different age-sex classes within each division (sex: males, females; age: juveniles (1month–4 years), subadults (4–10 years), young adults (10–18 years), prime-age adults (18–40 years), old adults (>40 years), Methods). Next, we obtained the monthly time-series of environmental predictors, i.e., total rainfall, average daily temperature at 2m above ground, and two aspects of primary productivity of vegetation (measured as remotely-sensed NDVI, Pettorelli et al. 2011): NDVI_div_, reflecting a division’s annual mean primary productivity, and NDVI_within-year_, i.e., monthly deviation from NDVI_div_ and thus reflecting seasonality in primary productivity (Methods). Furthermore, to explore potential survival consequences of these environmental predictors over a larger cumulative window of effects than the current focal month, we also recomputed the time-series as the rolling sums of rainfall and rolling averages of temperature and NDVI_within-year_, for up to four months including the current month (Methods). Our final dataset included observations for 43,257 division-months across age, sex categories and years. The raw average monthly mortality rate was 0.343% and an annualized mortality rate of 4.05%. Using this dataset, we developed 28 candidate models of monthly elephant mortality using generalized linear mixed effects models (GLMM, beta-binomial error structure, Methods), as described below.

### Baseline model

Our baseline GLMM accounted for the heterogeneity in monthly mortality risk across age, sex and division categories (fixed effects), while accounting for the random effects of current year and the temporal autocorrelation across months. The overall predicted monthly mortality risk was 0.22% across all age, sex and division classes (based on *emmeans*). Monthly mortality risk was higher in males (0.24%) than females (0.20%). Young adults (0.09%) and prime-age adults (0.11%) were least likely to die than juveniles (0.34%) and old adults (0.39%). There was also geographical variation in mortality across divisions (Fig. S5).

### The effects of rainfall, temperature and vegetation productivity on elephant mortality

Rainfall, temperature and NDVI showed large variation and were only weakly correlated in our dataset (Fig. 1, Pearson’s *r* <0.45). To evaluate the effect of these predictors, we examined several candidate models extending the baseline model by considering additive effects as well as the two-way interactions between temperature and rainfall, and between temperature and NDVI variables (Table S2, Methods). Based on preliminary inspections of non-linear effects (Methods, SI text b), we specified non-linear effects only for temperature (natural splines, *df*=2, Methods), whereas linear effects were retained for other predictors. To limit model complexity, we did not include interactions between environmental variables and categorical variables such as age, sex or division. Our focus was on inferring population-level responses, but we acknowledge that responses may vary across demographic and spatial groups.

From all 28 candidate models, we identified a *confidence set* of competing models within six AIC points of difference with the best-supported model (Harrison et al. 2018); these models accounted for over 98% total model weight and substantially outperformed the baseline model (Table 1). To make inferences that account for the uncertainty arising from model selection, we evaluated the consistency of estimated effects across the subset models containing each predictor, focusing on both the direction and magnitude of effects, rather than relying on a single best-fitting model. For all predictors, the direction of estimated marginal effects (i.e., effect at mean/reference values of other predictors) was largely consistent across models containing them, although their magnitude varied. Below, we report the marginal means of estimated changes in the odds of mortality obtained from these models summarized in Table 2 and Table S4–S7.

**Table 1.** The confidence set of top models of monthly mortality (ΔAIC ≤ 6.0 relative to top model). b = baseline model which includes the fixed effects of age, sex and division categories, and random effects of year and temporal autocorrelation across months (AIC=12216.41). r = rainfall, r_3_ = rainfall summed over three months, t = temperature (natural splines, *df*=2), n_d_ = annual mean NDVI of the division, n_w_= within-year centered NDVI. See Table S2 for AIC-based comparison and AIC_w_ of all models explored.

| Model | k | AIC | $\Delta AIC$ | $AIC_w$ | Model rank |
| --- | --- | --- | --- | --- | --- |
| $b + r_3 + (t \times n_d)$ | 27 | 12146.9 | 0.000 | 0.505 | 1 |
| $b + r_3 + t + n_d$ | 25 | 12149.24 | 2.334 | 0.157 | 2 |
| $b + (r_3 \times t) + (t \times n_d)$ | 29 | 12149.68 | 2.772 | 0.126 | 3 |
| $b + r_3 + t + n_d + n_w$ | 26 | 12150.95 | 4.046 | 0.067 | 4 |
| $b + (r_3 \times t) + (t \times n_d) + n_w$ | 30 | 12151.12 | 4.212 | 0.061 | 5 |
| $b + (r_3 \times t) + n_d$ | 27 | 12152.09 | 5.186 | 0.038 | 6 |
| $b + (r \times t) + (t \times n_d) + n_w$ | 30 | 12152.68 | 5.777 | 0.028 | 7 |

**Table 2.** Effect sizes expressed as odds of mortality for different fixed effects in the top-ranked model. The odds of mortality are with respect to mortality at the reference category for categorical predictors and at the mean value for continuous predictors.

| Effect | Odds of mortality | 95% CI (low) | 95% CI (high) | P |
| --- | --- | --- | --- | --- |
| Age class (ref: juvenile) | 1.0 | - | - | - |
| Age: subadult | 0.987 | 0.832 | 1.172 | 0.885 |
| Age: young adult | **0.267** | **0.214** | **0.333** | **<0.001** |
| Age: prime-aged adult | **0.327** | **0.279** | **0.385** | **<0.001** |
| Age: old adult | 1.136 | 0.968 | 1.333 | 0.117 |
| Sex (ref: female) | 1.0 | - | - | - |
| Sex: Male | **1.213** | **1.094** | **1.345** | **<0.001** |
| Division (Ref: Ayeyarwaddy) | 1.0 | - | - | - |
| Division: Bago East | 1.338 | 0.634 | 2.821 | 0.445 |
| DivBago West | 0.805 | 0.387 | 1.673 | 0.561 |
| DivChin | **2.995** | **1.308** | **6.855** | **0.009** |
| DivKachin | **4.009** | **1.819** | **8.834** | **0.001** |
| DivMagway | 1.063 | 0.527 | 2.141 | 0.865 |
| DivMandalay | 1.170 | 0.567 | 2.416 | 0.671 |
| DivNay Pyi Taw | 1.929 | 0.905 | 4.111 | 0.089 |
| DivRakhine | 2.065 | 0.935 | 4.561 | 0.073 |
| DivSagaing | 1.411 | 0.690 | 2.887 | 0.346 |
| DivShan North | **2.957** | **1.394** | **6.275** | **0.005** |
| DivShan South | 1.490 | 0.668 | 3.325 | 0.330 |
| Continuous predictor (Ref: mean) | 1.0 | - | - | - |
| 3-month rainfall (+1 SD) | **0.816** | **0.761** | **0.875** | **0.000** |
| NDVI<sub>div</sub>: +1SD (at mean temp*) | **0.746** | **0.602** | **0.925** | **0.004** |
| NDVI<sub>div</sub>: - 1SD (at mean temp*) | **1.340** | **1.081** | **1.661** | **0.004** |
| *Temp: +2 SD (at mean NDVI<sub>div</sub>) | 1.244 | 0.967 | 1.600 | 0.090 |
| *Temp: +1 SD (at mean NDVI<sub>div</sub>) | 1.082 | 0.986 | 1.188 | 0.094 |
| *Temp: -1 SD (at mean NDVI<sub>div</sub>) | 0.997 | 0.942 | 1.056 | 0.930 |
| *Temp: -2 SD (at mean NDVI<sub>div</sub>) | 1.054 | 0.901 | 1.232 | 0.512 |
| *Temp: +2 SD (at +1 SD NDVI<sub>div</sub>) | 0.950 | 0.629 | 1.434 | 0.806 |
| *Temp: +1 SD (at +1 SD NDVI<sub>div</sub>) | 0.994 | 0.852 | 1.158 | 0.934 |
| *Temp: -1 SD (at +1 SD NDVI<sub>div</sub>) | 0.958 | 0.876 | 1.047 | 0.341 |
| *Temp: -2 SD (at +1 SD NDVI<sub>div</sub>) | 0.883 | 0.694 | 1.124 | 0.313 |
| *Temp: +2 SD (at -1 SD NDVI<sub>div</sub>) | **1.628** | **1.234** | **2.149** | **0.001** |
| *Temp: +1 SD (at -1 SD NDVI<sub>div</sub>) | **1.179** | **1.067** | **1.303** | **0.001** |
| *Temp: -1 SD (at -1 SD NDVI<sub>div</sub>) | 1.039 | 0.959 | 1.126 | 0.350 |
| *Temp: -2 SD (at -1 SD NDVI<sub>div</sub>) | **1.257** | **1.004** | **1.574** | **0.046** |

Marginal effects for continuous predictors were obtained by generating predicted log-odds using *emmeans*, keeping other effects constant, and then exponentiating to obtain a marginal odds ratio (contrasts obtained using *revpairwise* method and *response* type). \*As temperature involves interactions and non-linear effects (spline terms), its predicted marginal effects for up to 2SD difference from mean temperature are shown at three discrete NDVI<sub>div</sub> values.

### Rainfall effects

Consistent with Prediction 1, elephant mortality strongly declined with rainfall in all models, with more support for the effect of 3-month cumulative rainfall than current-month rainfall (Table 1). An increase in 3-month cumulative rainfall by one standard deviation (SD=496 mm, relative to mean=501 mm) significantly reduced the odds of mortality by 17.5–20.4% (Figure 2, Table 1, Table S4-S6), similar to the 17.6% reduction in mortality associated with one SD increase in current-month rainfall (SD=191 mm, mean=168 mm) (Table S7).

**Figure 2.**
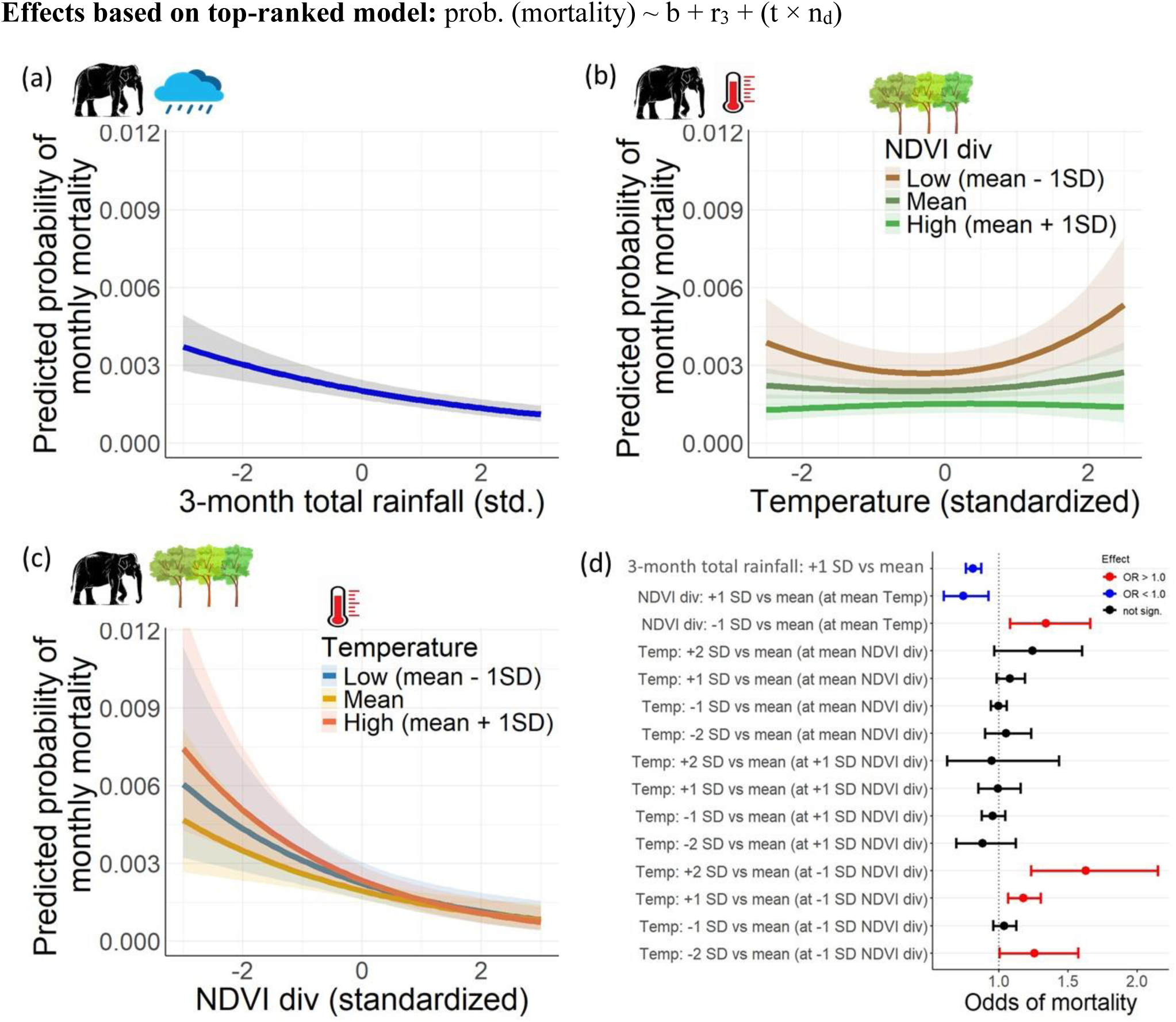
Effects based on the top-ranked model. Predicted mortality rates across standardized effects of a) 3-month cumulative rainfall, b) temperature (response shown for three discrete NDVI_div_ values due to interaction effects), and c) NDVI_div_, i.e., annual mean vegetation greenness in a division (response for three discrete temperature values); d) Changes in odds of mortality with respect to mean value for environmental predictors. Response curves were obtained using *emmeans* and are averaged across age, sex and division categories. b = baseline model terms described in Results.

### Temperature effects and their buffering by NDVI_div_

Consistent with Prediction 2, temperature showed well supported non-linear effects on mortality, with strong support for the magnitude of such effects to depend on NDVI_div_ and/or rainfall (Table 1, Table S4–S7). Mortality risk was higher at extreme temperatures especially when NDVI_div_ was low, with these U-shaped effects being asymmetric and marked by stronger effects of heat than cold (Figure 1). At low NDVI_div_, a one SD increase in temperature (i.e., at 26.87 °C vs. at mean=23.67 °C) increased the odds of mortality by 17.9–32.5% (*P*<0.05) whereas a one SD decrease in temperature (i.e., at 20.45 °C) had weak non-significant effects (3.9–19.6%, *P*>0.05) (Table 1, Tables S5, S7). Only more extreme cold temperatures (two SD below the mean or 17.23 °C) significantly increased mortality risk (by 25.7–101%) but similarly extreme hot temperatures markedly inflated the mortality risk (by 62.8–151%) at low NDVI_div_. Similar non-linear trends were also observed at mean NDVI_div_ but the effects were statistically not significant for up to two standard deviations from mean temperature in three out of four models containing the temperature × NDVI_div_ interaction (Table 2, Tables S5, S7). Notably, high NDVI_div_ completely buffered against such effects of temperature extremes, consistent with Prediction 5 (Fig. 2, Fig. S6). We found robust support for U-shaped effects of temperature as all top models containing spline effects with *df*=2 were supported well relative to their variant models where we altered the shape of temperature effects using splines (*df*=1–4, Table S3). However, the asymmetric shape of temperature effects inferred from its spline terms (*df*=2) received only marginally better support than models with symmetric quadratic effects (Table S3).

### Amplification of temperature effects by rainfall

The interactive effects of temperature and rainfall on elephant mortality contradicted Prediction 3, which expected low rainfall to intensify the effects of temperature extremes. Instead, an increase in rainfall, while enhancing survival at mean temperatures, amplified mortality under temperature extremes, particularly under hot conditions (Figure 3). However, the magnitude of such amplification depended on the time window considered for rainfall: the effects of heat were amplified more strongly by current-month rainfall than by rainfall summed over a 3-month window, which emphasizes immediate than delayed impacts (Figure 3). Relative to mortality at mean values of all predictors, a concurrent one SD increase in temperature and current-month rainfall increased the odds of mortality by 57.6% (Table S7) whereas this increase was 24.5–27% for 3-month rainfall (Table S5–S6); this trend was stronger at more extreme hot temperatures (Figure 3).

**Figure 3.**
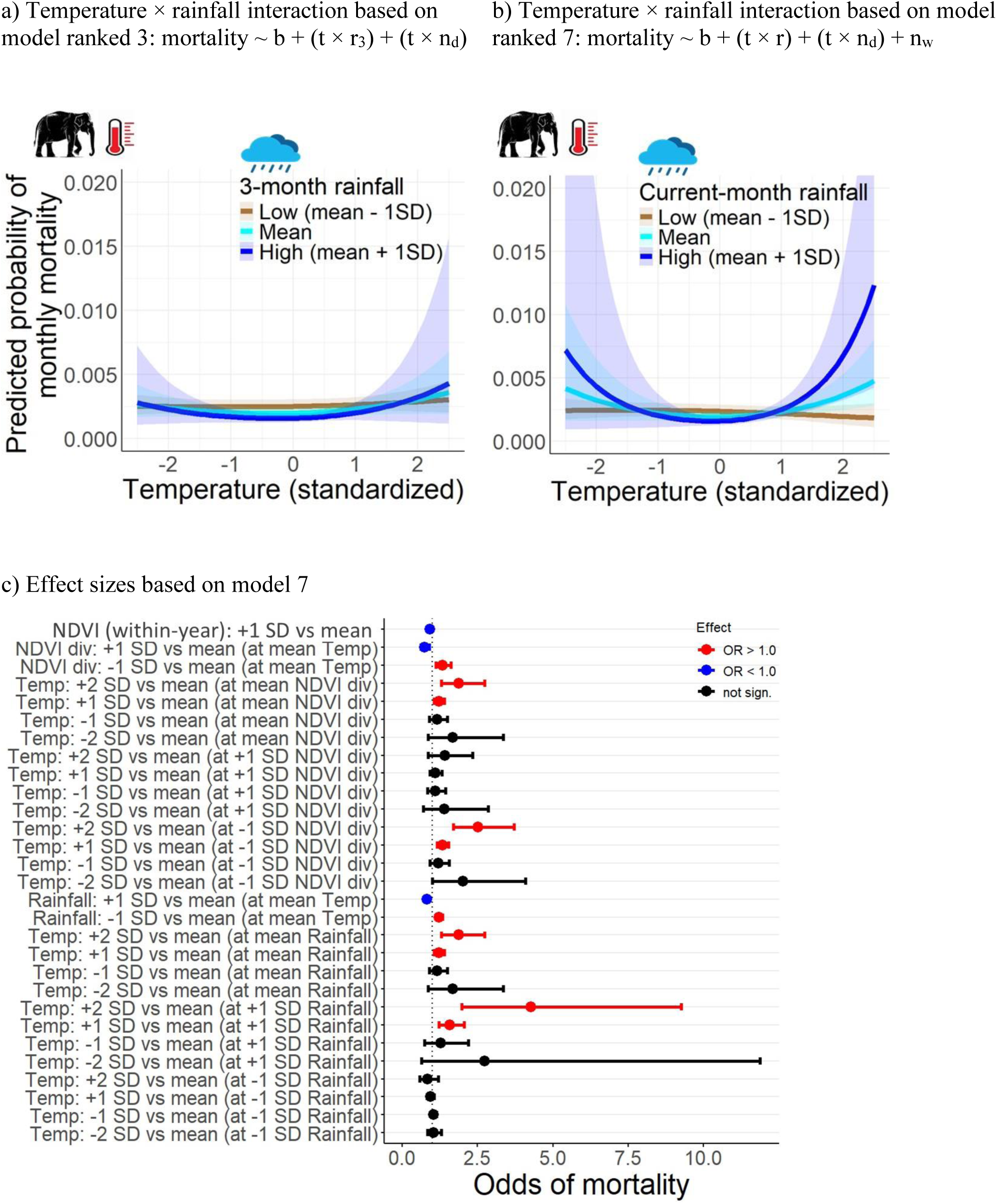
The interactive effects of temperature and rainfall on predicted elephant mortality rates based on a) temperature × 3-month rainfall interaction (from model ranked 3) and b) temperature × current-month rainfall interaction (from model ranked 7), and c) Effect sizes obtained from the model containing effects from current-month rainfall and within-year centered NDVI (model ranked 7). In panels a and b, predicted mortality rates across standardized values of temperature are shown for three discrete values of rainfall due to interaction effects. Response curves were obtained using *emmeans* and are averaged across age, sex and division categories.

### Effects of NDVI_div_ and NDVI_within-year_

All top models unanimously supported a buffering effect of NDVI_div_ (Prediction 4, Figure 2), with a one SD increase in NDVI_div_ associated with a 17.5–52.3% decline in the odds of mortality (Table 1, Table S4–S7). NDVI_div_ also strongly buffered against the effects of temperature extremes, as mentioned above. In contrast, we found weak support for the effect of vegetation seasonality (Prediction 6), as the odds of mortality increased by only 2–8.6% with a one SD increase in NDVI_within-year_ which captures the seasonal deviation from NDVI_div_ (Table S4–5, S7). Finally, we did not find support for Prediction 7 that the estimated effects of temperature extremes may be underpinned by seasonal drops in vegetation productivity. The presence or absence of NDVI_within-year_ in a model had low or negligible influence on the estimated effects of temperature extremes, as accounting for NDVI_within-year_ effects only marginally reduced the estimated effects of heat and marginally increased the estimated effects of cold extremes (Table S4–5, S7). In our dataset, months with lowest NDVI_within-year_ overlapped with temperature peaks (March–May) whereas the coldest months (December–February) marked the onset of decline in NDVI_within-year_ (Fig. S7).

## DISCUSSION

Our study fills an important gap in understanding the demographic impacts of climatic variation in animals with exceptionally large body size and slow life-histories. Mortality in this population of Asian elephants increased at temperature extremes, but such U-shaped effects of temperature depended on vegetation productivity and rainfall, underlining effects from multiple stressors. Our findings also suggest considerable bottom-up regulation, as high vegetation productivity of divisions not only reduced mortality but also strongly buffered elephants from the effects of temperature extremes. Since mortality is a key determinant of population growth, these findings highlight multiple pathways through which climate change can affect megaherbivore populations.

### The complex survival consequences of temperature regimes

Despite the recognition that global warming threatens endotherms, its demographic consequences in megaherbivores remain poorly known due to the scarcity of individual-based data (Boyles et al. 2011; Hetem et al. 2014; Fuller et al. 2016). Addressing this gap, we show that both extreme hot and cold temperatures negatively impacted survival in the Asian elephant, with such effects strongly evident at low vegetation productivity and high rainfall. To our knowledge, the giraffe is the only other megaherbivore where the survival consequences of temperature regimes were quantified using individual-based data, and it was found that their mortality was high in cold but not hot conditions (Bond et al. 2023). The tolerance of giraffes to heat may result from morphological features facilitating heat loss through increased body surface area (e.g., long neck and legs) and thus providing buffering against high heat stress (Bond et al. 2023). In contrast, high temperatures may impose greater heat stress in other megaherbivores whose “roundish” body outline does not provide optimal body surface-area/volume ratio for effective thermoregulation (e.g., all species of rhinos, elephants and hippos). In such animals with low surface-area/volume ratio, heat dissipation is slow and may be mostly effective during cool nights (Weissenböck et al. 2012). While elephants have large ears that help dissipate heat through increased surface area and high density of blood vessels (Dunkin et al. 2013), high mortality in hot conditions indicates that such functions of ears were insufficient to overcome heat stress. Therefore, other “round-bodied” megaherbivores, which have smaller ears than elephants, are expected to face similar or worse demographic impacts of temperature extremes. In our study population, while the seasonality of workload may certainly influence stress experienced by elephants (Mumby et al. 2015), it is noteworthy that the hot season coincides with the resting season (March–June), implying heat stress rather than workload as the underlying contributor to mortality in hot months.

Our analyses elucidate multiple aspects of the U-shaped effects of temperature found earlier based on a smaller dataset (Mumby et al. 2013). First, by expanding our analyses to a broader geography and environmental space (Fig. S2), we show that negative survival consequences of extreme heat and cold are stronger under low annual vegetation productivity of administrative divisions, whereas high vegetation productivity buffers against such effects (discussed below). Second, our findings suggest an asymmetric shape of temperature effects, with heat having stronger effects than cold. These findings point towards a stronger demographic signal in the hot range rather than the cold ranges of temperature regimes, although we also found considerable support for the symmetric effects of both heat and cold (Table S3) identified previously (Mumby et al. 2013). Third, we ruled out seasonal drops in vegetation productivity as an underlying explanation for the effects of temperature extremes, which remained strong even after controlling for vegetation seasonality (Figure S6, Table S4–5, S7). In fact, temperature was only weakly correlated with vegetation seasonality measured as NDVI_within-year_ which declined only moderately in the coldest months (December–February). However, hot months coincided with the largest drop in NDVI_within-year_ (Fig. S6) which may explain its weak influence on the estimated effects of heat (Table S4, S7). Finally, contrary to our expectation that low rainfall would intensify the effects of heat on mortality, we instead found intensified effects of heat at high rainfall (Figure 3), suggesting that high humidity amplifies heat stress. This appears consistent with elevated fecal glucocorticoid concentrations in a subset of this population during June–August when both heat and rainfall are high (but also coinciding with onset of work season, Mumby et al. 2015), which highlights the need to directly assess the link between physiological responses and heat-humidity stress.

Multiple biological mechanisms could have generated such U-shaped effects of extreme temperatures and their amplification by high rainfall. First, inflated mortality in months with concurrently high rainfall and temperature could arise from the limits placed by high humidity on dissipation of heat via evaporation. Negative impacts of concurrently hot and humid conditions are often seen in animals, including humans where few hours of exposure to “wet-bulb temperatures” above 35 °C can be lethal because high humidity constrains heat dissipation through evaporation and sweating (Raymond et al. 2020; Coulson et al. 2025). Such lethal thresholds of wet-bulb temperature may possibly be lower in megaherbivores whose skin has very few sweat glands and thus can dissipate heat largely through pores (eg., all species of elephants and rhinos, but not giraffes). Second, while behavioural adjustments like seeking shade, mud-bathing and sprinkling water on body can ease thermoregulation, these behaviours reduce foraging opportunities (Mole et al. 2016; Baskaran et al. 2010). The resulting starvation can limit thermoregulatory abilities (Rey et al. 2017), potentially promoting a costly spiral of starvation and body heating impacts. As larger animals have greater food requirements and foraging time (Owen-Smith 1988), how climate effects vary along the body size continuum is an interesting arena for future investigations. Third, mortality in cold conditions could result from the physiological costs of navigating cold stress (Khaliq et al. 2014). Minimum temperatures in Myanmar dropped to below 12 °C in several years (averages in Figure S6, and often below 0 °C in some places) which can generate cold stress. With increasing body size, refuges to escape cold become non-existent (unlike burrows available to small mammals), whereas huddling together for warmth for prolonged hours comes with foraging costs. Incidentally, deaths from “general weakness” as diagnosed by veterinarians are more likely under cold conditions in this population (Mumby et al. 2013). Furthermore, in our study, higher mortality in cold conditions was mostly evident at low annual vegetation productivity. Fourth, extreme temperatures may also promote disease-linked mortality through proliferation of pathogens, parasites and their vectors adapted to such extremes (Cohen et al. 2018; Nuttall 2022). In this population, mortality linked with infectious diseases is more common in hot and wet months whereas deaths linked with non-infectious diseases occurred more in cold months (Mumby et al. 2013). Ultimately, the emergent shape of the temperature-mortality curve may depend on the cumulative effects of distinct causes of deaths.

### High vegetation productivity buffers megaherbivore populations

Consistent with the strong bottom-up limitation expected for megaherbivores, we found that elephant survival was substantially better in divisions with high annual mean NDVI, which also provided a buffer against the negative impacts of extreme temperatures. This buffering observed in high-NDVI divisions likely reflects enhanced habitat suitability (e.g., in Sumatra: Rood et al. 2010) which may arise from multiple vegetation attributes. NDVI could partly reflect the productivity or diversity of elephant food plants, but only ground-truthing can verify this relationship at such large scales. For instance, NDVI did not reflect the abundance of common food species at very fine scales in southern India’s forests (Gautam et al. 2019). However, that study assumed that grass dominates elephant diet, whereas later work found elephants in the same forests to predominantly feed on browse (Gautam et al. 2025) which may be better captured by NDVI. Alternately, better survival in high-NDVI habitats could also arise from microclimatic buffering provided by dense canopy shade against heat spikes (Santos et al. 2026). However, since NDVI may also be correlated with forest management practices and other differences among divisions, the mechanisms underlying improved survival in high-NDVI habitats require further investigation.

While vegetation seasonality, captured as NDVI_within-year_, weakly improved survival, it was not the underlying explanation of high mortality at temperature extreme. These effects of NDVI seasonality may have been limited due to its failure to capture other seasonal aspects of vegetation that are important for elephants. For example, NDVI may have poorly captured forage quality. Like in other herbivores (Drescher et al. 2006), it is possible that elephant forage also shows seasonal maturation that accompanies build-up of less nutritional tissues and chemical defenses. In such scenarios, peak forage quality may coincide with intermediate rather than peak vegetation greenness. Incidentally, crude protein content in the common food plants (and in milk) of a subset of elephants from our study population was low during December–February, a period when NDVI is not the lowest (Fig. S4) relative to July–Sept in monsoon when forage quality was better (Dierenfeld et al. 2020) but NDVI was only moderately high.

### Threats and silver linings in a changing world

Studies on animal life-histories emphasize that demographic impacts of environmental disruptions generally vary along the slow-fast continuum, with the populations of short-lived species typically being more sensitive whereas long-lived species appear more buffered against short-term perturbations (Morris et al. 2008; Jackson et al. 2022). While we did not quantify population growth rates, our findings based on individual-level data at a very fine temporal resolution demonstrate detectable demographic consequences of environmental variation, consistent with findings on other long-lived species (Bond et al. 2023; Lee et al. 2011; Ndlovu et al. 2023). These mortality responses are particularly relevant for modelling population persistence of megaherbivores whose low reproductive rates imply slow recovery after demographic disruptions.

The empirical patterns emerging from this large population inform predictions on how long-lived, large-bodied animals may respond to climate change. First, the U-shaped effects of temperature imply that global warming would elicit divergent demographic responses in hot vs. cold conditions. While a shift towards more extreme heat would increase mortality during summers, survival in cold periods may improve as warming may shift populations to the less steep parts of the cold-response curve, possibly offsetting some deaths from hot months. However, the net demographic impacts of global warming may still be unfavourable, because i) mortality itself increased more in response to heat than cold in our study, ii) night-time temperatures, which generally facilitate loss of body heat gained during the day (Weissenböck et al. 2012; Dunkin et al. 2013), are expected to increase more than day-time temperatures (Anderegg et al. 2015), and iii) these tropics are projected to face intensification of extreme heat whereas extreme cold may become dampened (Kodra and Ganguly 2014). Second, mortality risk arising from concurrently hot and humid conditions may increase because extreme heat and precipitation events in the tropics are projected to increase by over 70% (Tang et al. 2025). Such risks from concurrent heat and humidity may be greater in hot-and-humid habitats than hot-and-dry habitats; incidentally, such effects are not seen in individual-based studies from dry habitats of Africa (Bond et al. 2023; Thorley et al. 2025). Third, an interesting plausibility is the projected long-term rise in vegetation productivity in the tropics (as photosynthetic performance by plants may increase due to warming and CO_2_ fertilization, Nemani et al. 2003; Pan et al. 2014). This could be another silver lining for large herbivores if canopy shade and forage production increase with vegetation productivity, assuming that existing habitats are not lost to human activities. Fourth, more frequent pulses of herbivore mortality may result from greater frequency of droughts associated with increasingly erratic rainfall patterns in this region (Roxy et al. 2017). Finally, our findings of the deadly consequences of concurrent effects emphasize future challenges for megaherbivores, as climate change may create challenging local niches with multiple stressors.

In summary, our findings demonstrate that megaherbivore populations are regulated by the concomitant effects of vegetation productivity, rainfall and temperature regimes. Given the broad spatial extent and fine temporal resolution of our analyses, similar responses may be common in hot and humid habitats of Asia that support diverse but understudied assemblages of large herbivores. Our findings can help in identifying conditions posing demographic risks under climate change, informing both the management of semi-captive elephants and future rewilding efforts (Thitaram et al. 2024). Finally, long-term demographic data from wild populations remain essential to evaluate the generality of our findings, although demographic datasets with comparable spatiotemporal extent, resolution and bioclimatic breadth are extremely difficult to obtain for wild elephants, which highlights the unique value of this population for understanding climate-linked threats. Future comparisons between wild and semi-captive elephants inhabiting comparable bioclimatic niches could further clarify the extent to which our findings are generalizable.

## MATERIALS AND METHODS

### Demographic data

With exceptionally large body size, slow life history and long lifespan, the Asian elephant is an excellent model species to understand how long-lived megafauna are affected by climate change. We used the individual-based demographic records from Myanmar’s timber elephant population (Mar et al. 2012; Lahdenperä et al. 2018). This population occupies habitats ranging from evergreen, moist deciduous, dry deciduous and mixed deciduous, as well as savanna woodlands and scrub habitats (land-cover classes based on data from Myanmar Information Management Unit: https://themimu.info), thus encompassing a large diversity of hot and humid habitats where Asian elephant populations are found and for which climate effects on megaherbivores are poorly understood. Ages are accurately known for individuals born in captivity that represent two-thirds of all individuals in our final dataset (called captive-born, N=2837), whereas ages were estimated by MTE officials at the time of capture for the remaining one-third elephants that were captured from the wild (called wild-caught, N=1620) to sustain this population, although wild capture is illegal in Myanmar since 1990s. We extracted demographic data for the study period January 1982–December 2020 for all individuals which could be assigned to one of Myanmar’s administrative divisions or states (Figure S1) and excluded all individuals that were known to be translocated across divisions. We chose January 1982 as the start of our study period because before 1982 full-year monthly series were not available for a key predictor, NDVI (see below). We applied left-censoring to months before January 1982 (for all individuals born before and alive then) whereas right-censoring was done to exclude the months after the individual disappeared from the dataset due to death or was no longer tracked.

We then created a monthly time-series of each individual’s mortality status (1/0) while updating their age in subsequent months. The ages were binned into five exclusive age classes defined *a priori* to reflect the distinct life stages of these elephants: juveniles (1 month to 4 years when the individual is usually separated from the mother for taming/training), subadults (4–10 years, the approximate age when females can give birth), young non-working adults (10–18 years after which individuals join the workforce), prime-age working adults (18–40 years) and old adults (>40 years). We excluded neonates (<1month) because their mortality depends heavily on condition at birth and maternal factors (condition during gestation and lactation) than to the environment and is higher than other classes by an order of magnitude. Finally, using these time-series of all individuals within each administrative division, we obtained a composite time series of age-specific monthly mortality, specified as the total number of deaths and total number of individuals of each age-sex class present in each division during each month, spanning January 1982–December 2020. We excluded Tanintharyi division because of the poor quality of data for a key predictor, NDVI (see below; Tanintharyi also had smallest sample size). This space/time-based filtering resulted in a dataset with 4,457 individuals with known sex, age and division location for the whole study period.

### Climate and vegetation productivity data

We obtained climate and vegetation data for spatial extents where these elephants are managed by MTE within each of the 12 administrative divisions represented in the filtered demographic dataset. To only retain areas within the distribution range of these elephants, we excluded from each division’s polygon all the townships where MTE does not operate, as well as areas under agricultural land-use (division and township *.shp* files obtained from https://themimu.info). Matching the division-months in demographic data, we then extracted the monthly time-series for total rainfall (CHIRPS, Funk et al. 2015) and average daily temperature at 2m (ERA5 LAND daily aggregates, Muñoz-Sabater et al. 2021), using Google Earth Engine (Gorelick et al. 2017). These rainfall and temperature data products proved reliable as we found a good agreement with weather station data in four townships (Pearson’s *r* = 0.90 for temperature and 0.81 for rainfall, SI text a). These time-series were obtained as averages across pixels in each division. As a proxy of primary productivity of vegetation, we obtained the NDVI, a vegetation index widely used in animal ecology (Pettorelli et al. 2011). To extract the time-series for NDVI spanning the full study period, we sourced satellite remote sensing data from MODIS (MOD13A1 16-day composite product, 2000 onwards) and AVHRR (daily NDVI, before 2000) at a common resampled resolution of 1 km^2^. AVHRR images are available at daily resolution but images on some days can be of poor-quality due to clouds, haze, etc., whereas MOD13A1 uses best-pixel compositing strategy based on quality-controlled images to minimize such effects. To apply such a best-pixel compositing strategy to AVHRR data, we selected images after applying quality-assurance masking (high-quality, snow-free, cloud-free filters) and then extracted monthly maximum NDVI for each pixel within each month. This approach proved valid since it yielded a strong positive correlation between processed AVHRR and MODIS NDVI raw measurements for the months when both sources were available (overall Pearson’s *r* =0.84; Tanintharyi division was excluded as it showed poor agreement between these two datasets, scatterplots in Figure S3). We then obtained the two time-series of mean NDVI (across all pixels of each division) per month for the full study period. We developed the final NDVI time-series using MODIS data whenever available and AVHRR data otherwise.

Since we wanted to separate the effects arising from the seasonality of vegetation and the annual mean NDVI of the division, we employed within-subject centering approach (van de Pol and Wright 2009) to decompose monthly NDVI series into two distinct time series: a) NDVI_div_ i.e., a division’s annual mean NDVI (that remains same for all months in a year) and b) NDVI_within-year_ or within-year centred NDVI within each division, reflecting monthly deviation from NDVI_div_. All the four environmental variables were then scaled (with mean = 0 and SD =1) to make their effect sizes comparable in statistical models described below.

### Statistical analyses

We used these demographic and environmental data to build GLMMs with binomial or beta-binomial error structure using the R package *glmmTMB* (Brooks et al. 2017). Our main response variable was mortality (binomial outcome) and modeled as a two-column matrix, i.e., *cbind* (deaths, total cases - deaths), where deaths were the statistical successes and the remaining cases are failures (or survivals) across the binomial trials in each division-month. Our final dataset consisted of N=43,257 division-months, after applying abovementioned filters and excluding months for which we lacked data on either temperature, rainfall or NDVI. We first developed a *baseline model* which served as a base for further evaluating the effects of environmental predictors. The baseline model included the fixed effects of age class (levels: juveniles, subadults, young non-working adults, working adults and old adults), sex (male/female), and administrative division (12 divisions). The model also included a random effect accounting for the non-independence between years, as well as temporally auto-correlated random effect accounting for the non-independence between consecutive year-month observations within the same administrative division (using a first-order autoregressive or *AR1* correlation structure). To evaluate the structure of residuals, we inspected plots of simulated residuals using the *DHARMa* package (Hartig and Hartig 2017). We initially considered a binomial error structure, but since the binomial model showed signs of over-dispersion, we relied on a beta-binomial error structure instead.

We examined several *candidate models* expanding on this baseline model, to evaluate the effects of environmental predictors. Before this, we looked for any overt non-linear relationship between mortality risk and the quantitative proxies used for climate and vegetation. We did so by first converting the quantitative predictors into qualitative ones using original quintiles as levels, and then visually examining prediction plots obtained for each predictor added to the baseline model separately (SI text b). These plots suggested that only temperature had an overt U-shaped effect, with higher mortality at both extreme quintiles than at the middle quintile (Fig. S4). Therefore, we explored non-linear representations for modelling the effect of temperature, whereas linear effects were maintained for rainfall and NDVI. We modelled the shape of the effect of temperature using four forms of natural splines (*df*=1–4, representing linear to increasingly complex curves) and retained the parameterisation involving two degrees of freedom since it resulted in the lowest AIC (SI text b). Furthermore, we also examined if these variables had effects over cumulative time windows up to four months (including the current month), and found that only rainfall had cumulative window effects (three-month period, based on AIC values) better than those from current-month measurements (SI text c, Table S1). Therefore, while developing candidate models, we explored the effects from both three-month total rainfall and current-month rainfall, whereas we used only current-month measurements for temperature and NDVI. In these models, we considered similar effects across all age, sex and division classes although we acknowledge that the effects of environmental variables may be more nuanced across these demographic/spatial groups.

We developed 28 candidate models (Table S2) to understand the environmental predictors of mortality. We started by adding the additive effects of rainfall and temperature to the baseline model. Next, we added NDVI_div_ to account for the effects of a division’s mean annual primary productivity, and NDVI_within-year_ to account for the seasonal deviations from this annual mean. Next, we examined the potential effects of temperature interacting with rainfall and NDVI variables, to examine if extreme temperatures worsened the effects of dry conditions or dry vegetation. These included: a) rainfall × temperature interaction as rainfall could alter the effects of temperature, b) temperature × NDVI_div_ as divisions with higher vegetation productivity could buffer elephants against the effects of heat, and c) temperature × NDVI_within-year_ since seasonal drops in vegetation productivity could intensify the effects of temperature. We relied on model comparison to draw inference and retained a *confidence set* of top models within six AIC points of the best model (Harrison et al. 2018). To evaluate if a focal predictor had a “meaningful” effect, we relied both on the effect size by converting the estimated logit-odds converted to odds ratios (marginal effects). We evaluated the consistency of estimated effects across the subset models containing each predictor, focusing on both the direction and magnitude of effects, rather than relying on a single best-fitting model.

Finally, for each of these top models, we again inspected the shape of temperature effects by altering the degrees of freedom of natural splines (*df*=1–4, SI text b), as well as examining quadratic effects of temperature representing symmetric effects of heat and cold reported earlier (Mumby et al. 2013). We relied on the parametrization with the lowest AIC (Table S3).

## ACKNOWLEDGMENTS

We thank the Ministry of Natural Resources and Environmental Conservation in Myanmar, for their support and permission to work with the Myanma Timber Enterprise (MTE), and all the veterinarians and officials involved in data collection, especially Dr. Htoo Htoo Aung and Dr U. Kyaw Nyein. We also thank the Myanmar Timber Elephant Project members for data curation and compilation, and particularly Dr. Khin Than Win, Thuzar Thwin and Mu Mu Thein for inputs on locating MTE agencies in administrative divisions and for project coordination. This study was financially supported by the European Research Council project *KinSocieties* (to V.L., ERC-2022-ADG, grant number 101098266).

## Notes

Competing Interest Statement: Authors do not have any competing interests.

### Competing Interest Statement

The authors have declared no competing interest.

