## Supplementary material for "Megaherbivore mortality risk from temperature extremes is buffered by vegetation productivity but amplified by rainfall": SI

**This SI file includes:**

Supporting text a to c

Figures S1 to S7

Tables S1 to S7

SI References

Supporting Information Text

***a) Reliability of rainfall and temperature data***

Ground-based weather station data from four townships suggest that the monthly total rainfall (CHIRPS) and monthly average daily temperature at 2m (ERA5 LAND daily aggregates) derived using Google Earth Engine proved to be reliable (average Pearson’s r = 0.90 for temperature and 0.81 for rainfall). We found that the ground data based from the Gangaw, Katha, Mawlaik and Shwebo weather stations (source: Mumby et al. 2013) agreed well with temperature data based on ERA5 LAND (Pearson’s *r* = 0.93, 0.88, 0.93, 0.90) and the rainfall data based on CHIRPS product (Pearson’s *r* = 0.77, 0. 84, 0.85, 0.79) for polygons of townships with these names. Note that these polygons represent the entire township within which the weather station represents just a point, which could have affected the correlation.

***b) Non-linearity of environmental effects***

We considered non-linearity (at the scale of the linear predictor, i.e., logit scale) only for the effects of temperature, based on initial plot-based visual inspection of whether mortality in the middle quantiles of these predictors differed from both extreme quintiles (quintile 1 and 5, Figure S4). These inspections were based on simple models where only the variable of interest (modified as a qualitative variable classified into quintiles) was added to the baseline model (i.e., to model *b* in Table 1 in main text). Subsequently, we modelled the shape of the effect of temperature using four forms of natural splines with *df* = 1 to 4, where *df*=1 represents a linear effect and higher *df*s allow increasingly complex curvatures. To develop final candidate models listed in Table 1, we chose *df*=*2* after considering AIC-based comparisons of models since models with *df*=2 performed better than the linear model (ΔAIC > 6.0) and was the most parsimonious specification among other forms that were more complex but not substantially better. The AIC values for the four model variants were: 12218.36 (*df*=1), 12192.48 (*df*=2), 12194.37 (*df*=3), 12192.21 (*df*=4).

For our confidence set of seven top models (Table 1 in the main text), we again examined the curvature of temperature effects by comparing models with *df*=1 to 4 for the spline effect of the temperature term and found good support for these top models with *df*=2. These models with *df*=2, implying asymmetric U-shaped effects of temperature (heat having greater effect than cold, main text), were either better than more complex curvatures or were as competitive in terms of AIC values in six of the seven cases (Table S3). We further compared the asymmetric U-shaped effects of temperature in these models (*df*=2) with quadratic effects where both cold and heat have symmetric effects on elephant mortality. Model comparison suggests only marginally better support for models with asymmetric effects than symmetric effects (Table S3). Therefore, our findings indicate greater certainty in temperature effects where heat has greater impact than cold, although equal effects of heat and cold is also a plausible scenario with good support. In contrast, we find poorer support for only linear effects (models with *df=1*) where mortality increases linearly as temperature increases.

***c) Window of delayed effects***

In the main text, in addition to the linear effects of rainfall measured in the current month, our main analyses also examined models with rolling sums of rainfall over 3 months. This is because considering rolling sums over 3 months led to GLMMs with a better goodness of fit (as measured by AIC) than models considering the effect of rainfall in the current month, or summed over two months only (ΔAIC>6.0). Rolling sums over four months performed similarly well than rolling sums over three months (ΔAIC <1.0), but we retained the latter parameterization for the sake of parsimony. In contrast, the rolling effects over windows wider than a single month did not perform better than measurements from the current month for temperature and NDVI_within-year_. AIC values of different windows are shown in Table S1.

Figures

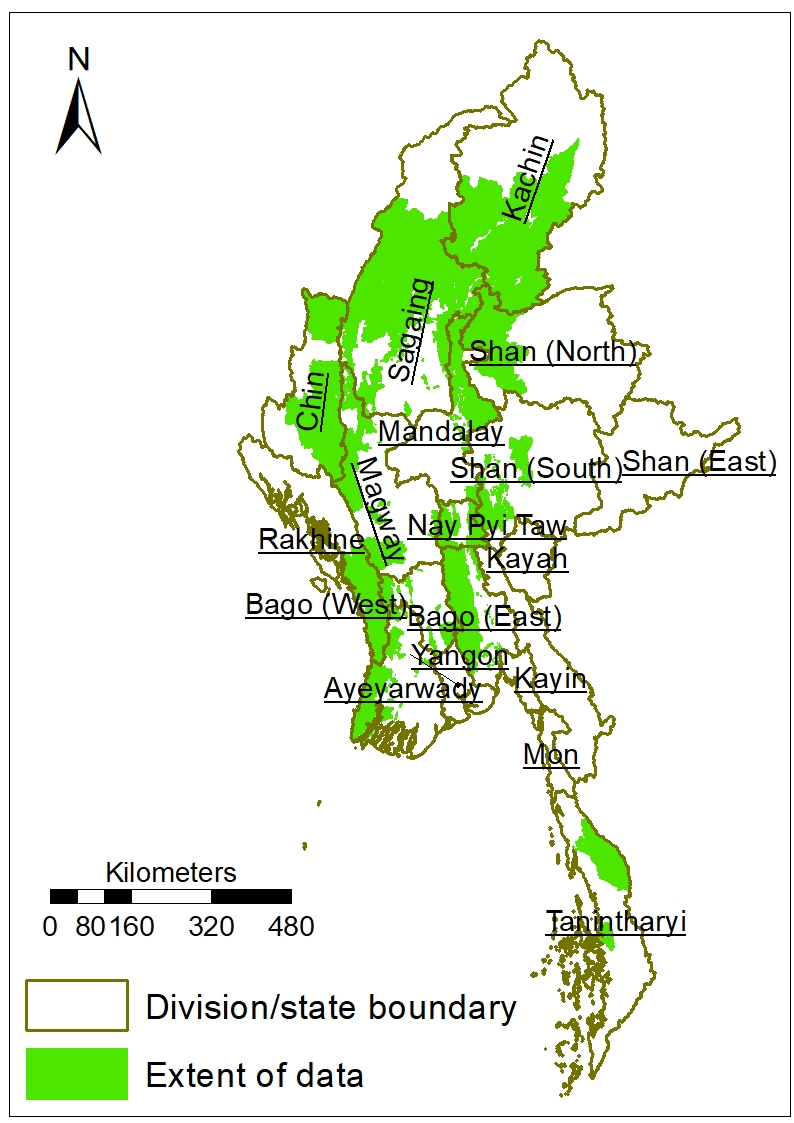

Fig. S1. Map showing the spatial coverage of MTE elephants considered in the study (in green). Tanintharyi division (southernmost) was excluded from the analyses due to poor quality of NDVI times series (due to poor agreement between two sources of data, i.e., MODIS and AVHRR, Fig. S2).

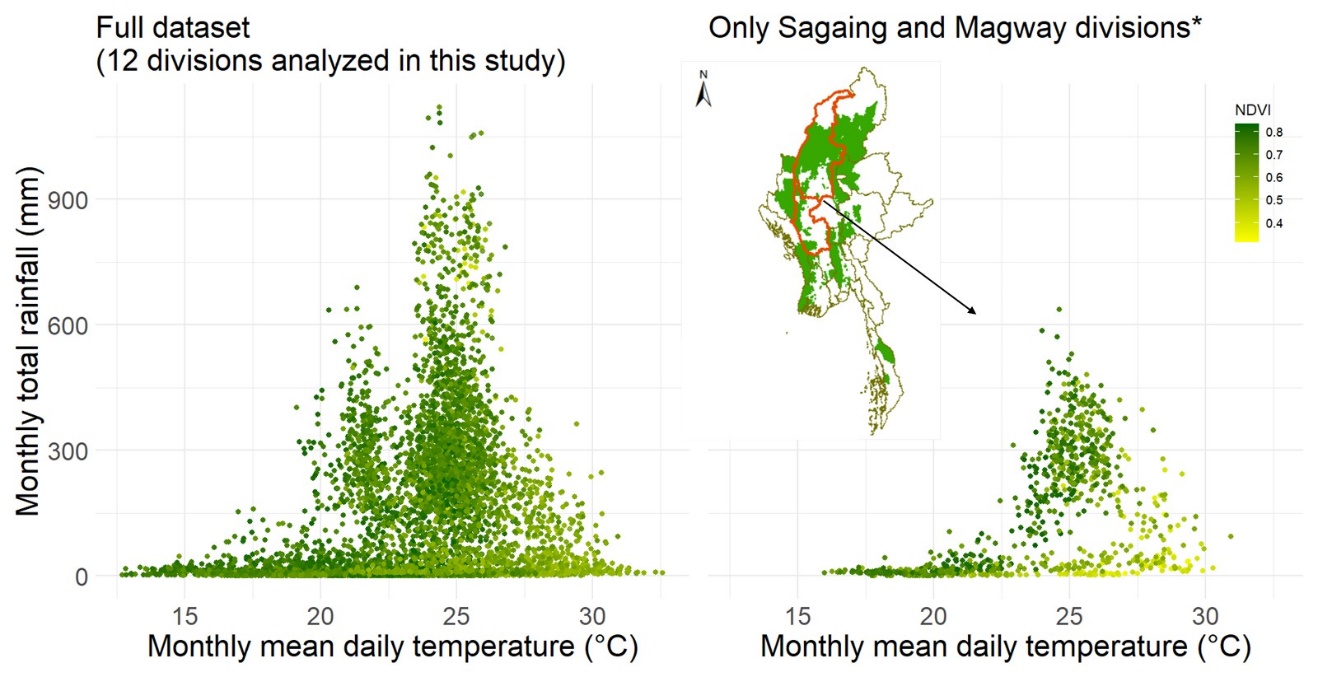

**Fig. S2**. The environmental width of the 12 administrative divisions covered in this study (left) and the two administrative divisions covered in a previous study on mortality in this population (right, *Mumby et al. 2013). The present study covers a much broader environmental niche.

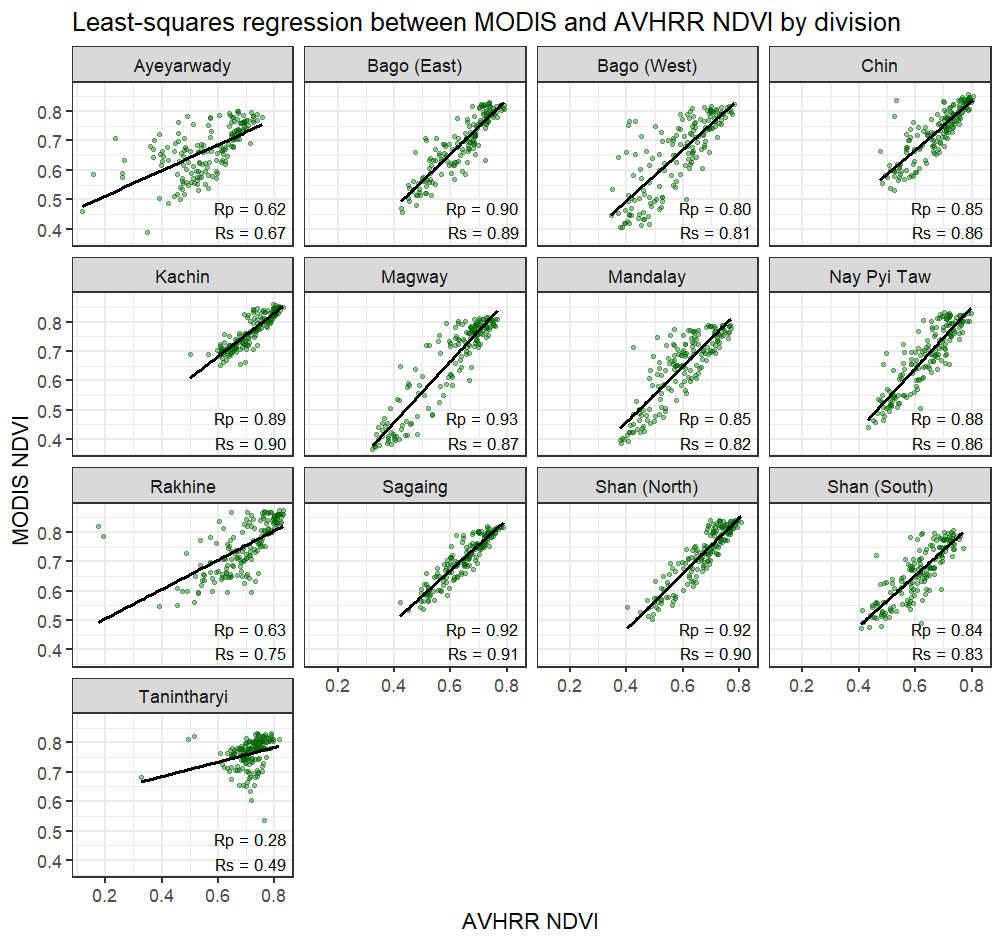

Fig. S3. Correlation between NDVI data derived from MODIS and AVHRR products within different divisions for 166 months for which quality-controlled images from both sources were available. Rp and Rs represent Pearson’s and Spearman’s correlation coefficients. Tanintharyi division was excluded from final demographic analyses because it showed poor agreement between these two datasets (possibly because its several small islands were adjacent to water that affects NDVI value of those pixels; map in Fig S1).

| a)  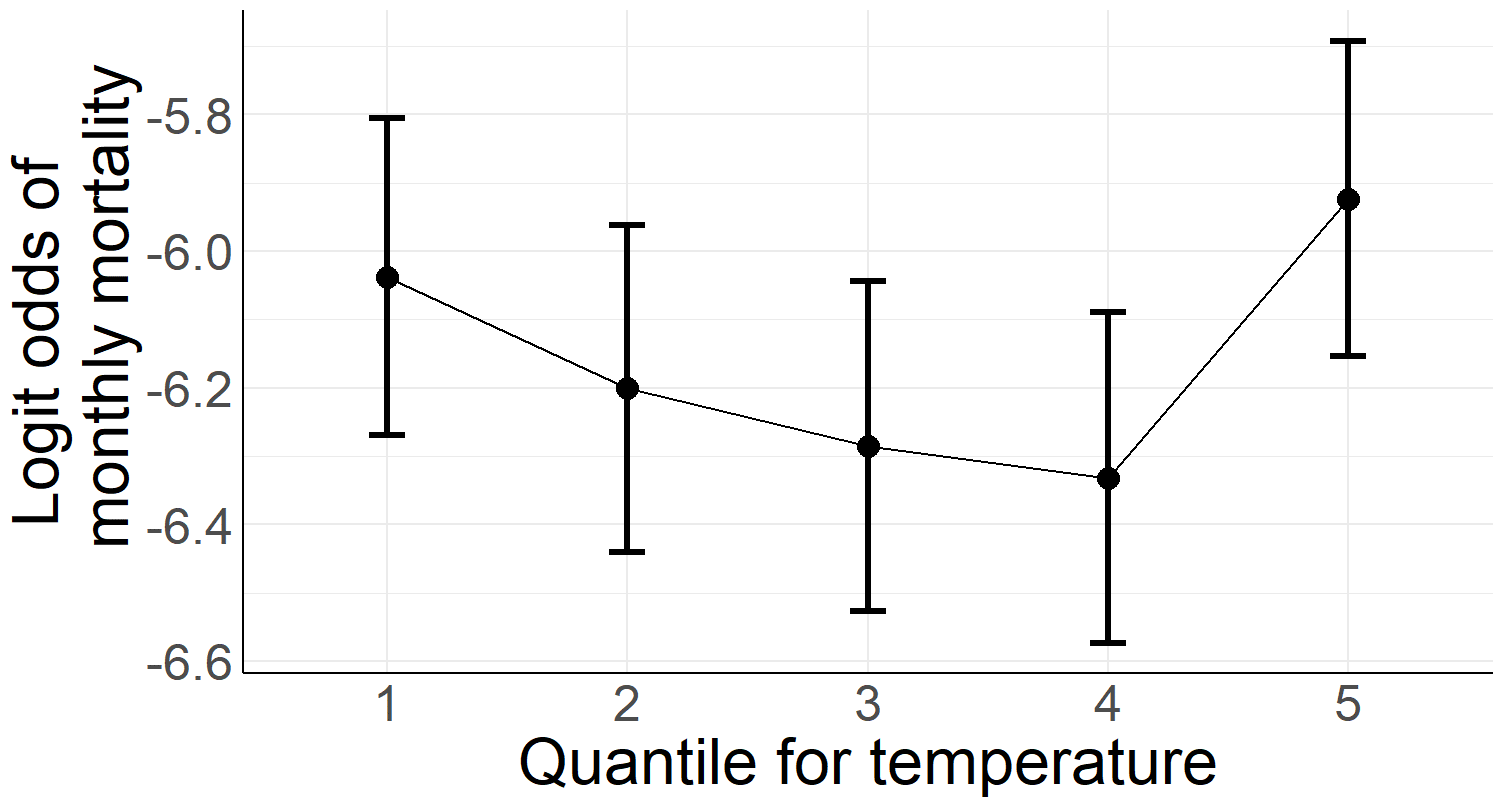 | b)  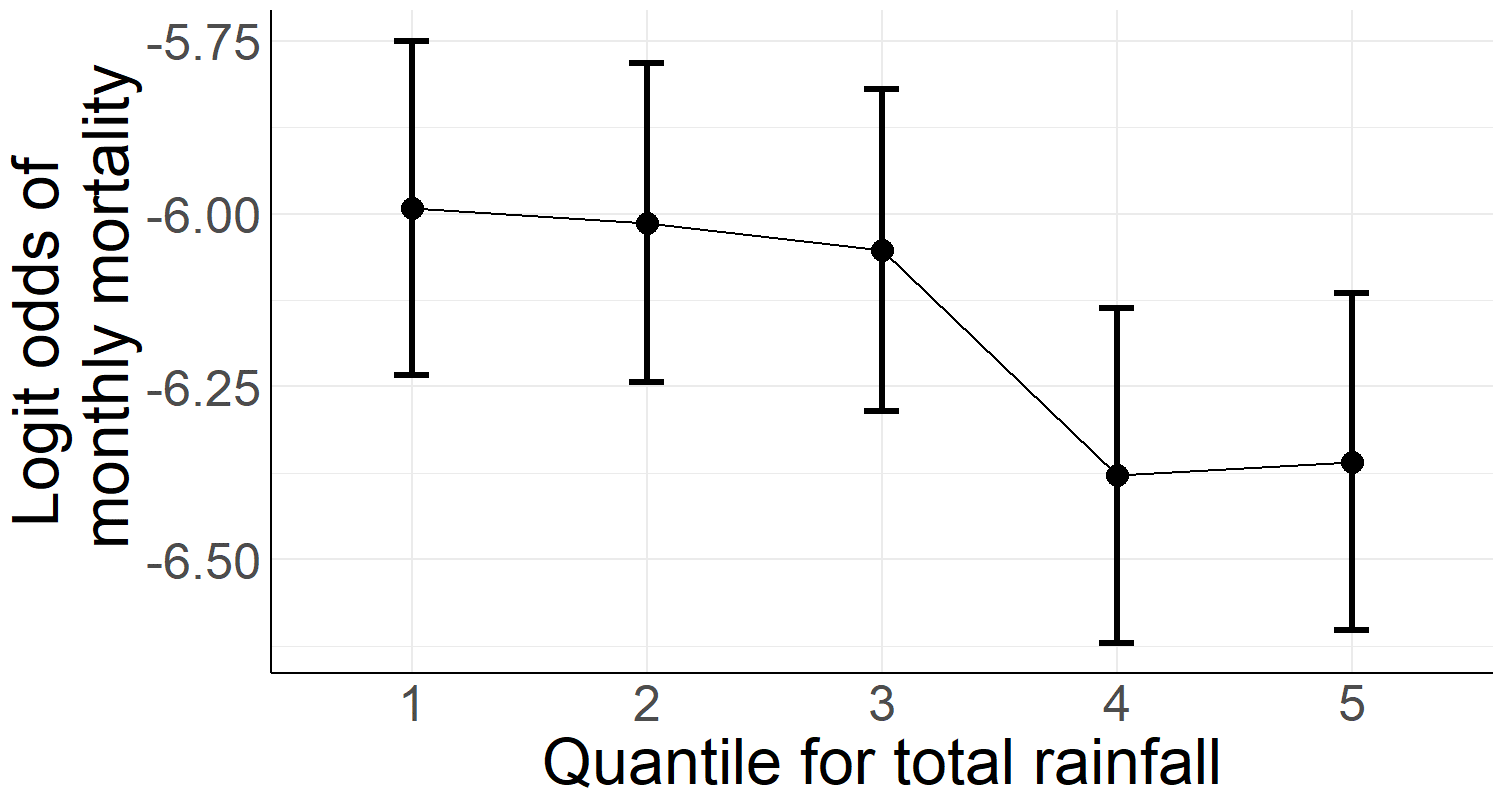 |
| --- | --- |
| c)  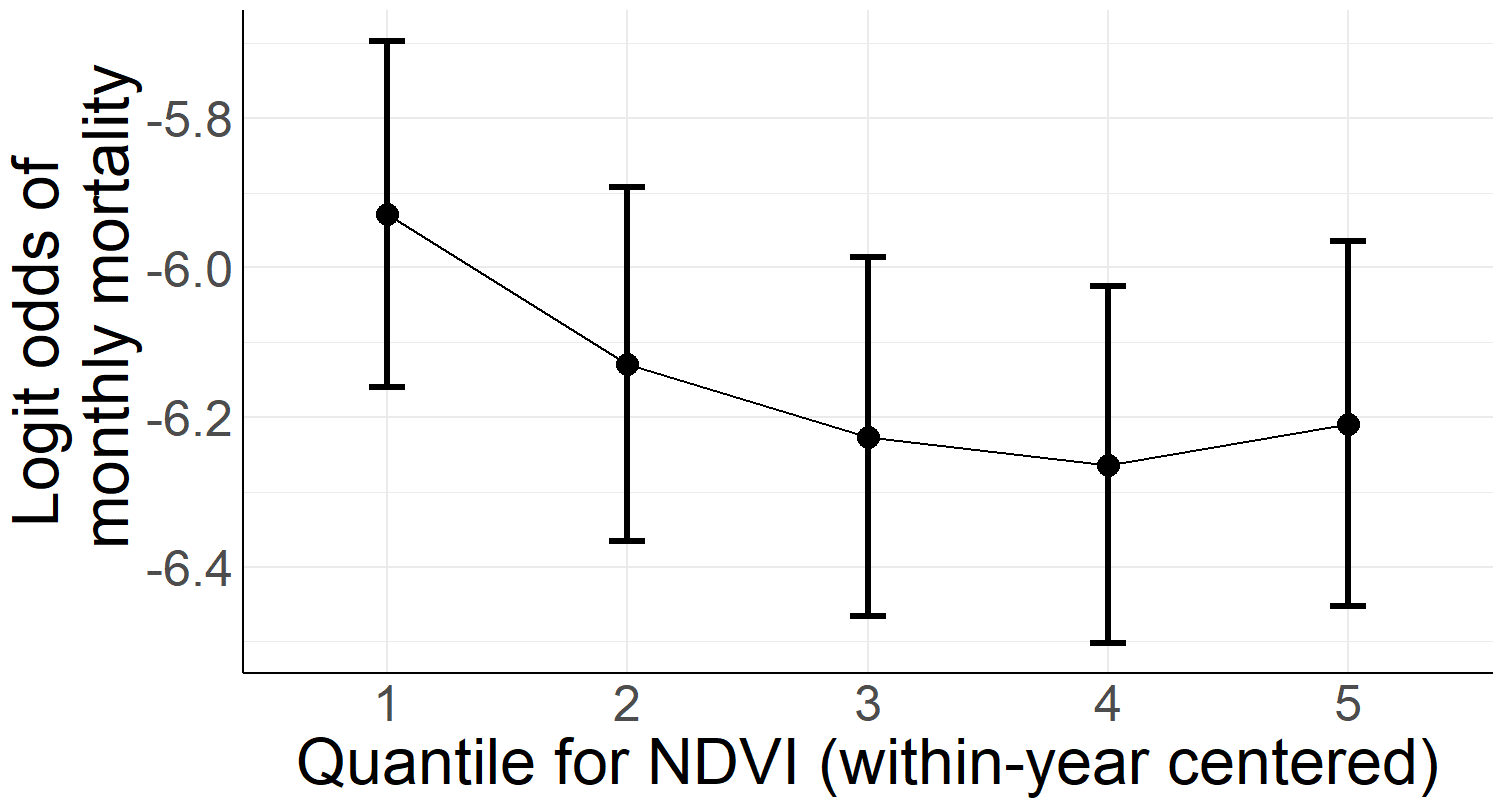 | d)  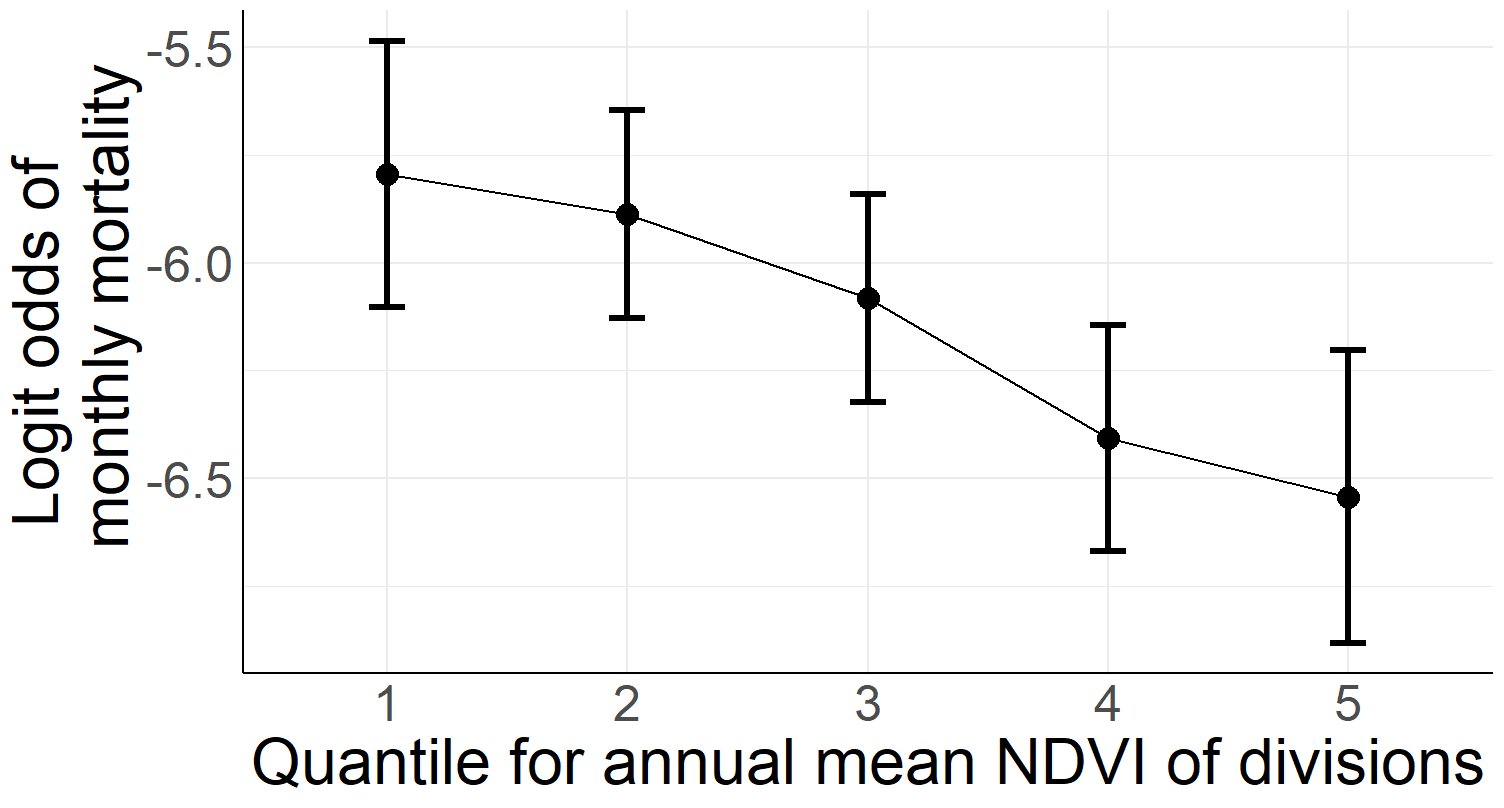 |
| e)  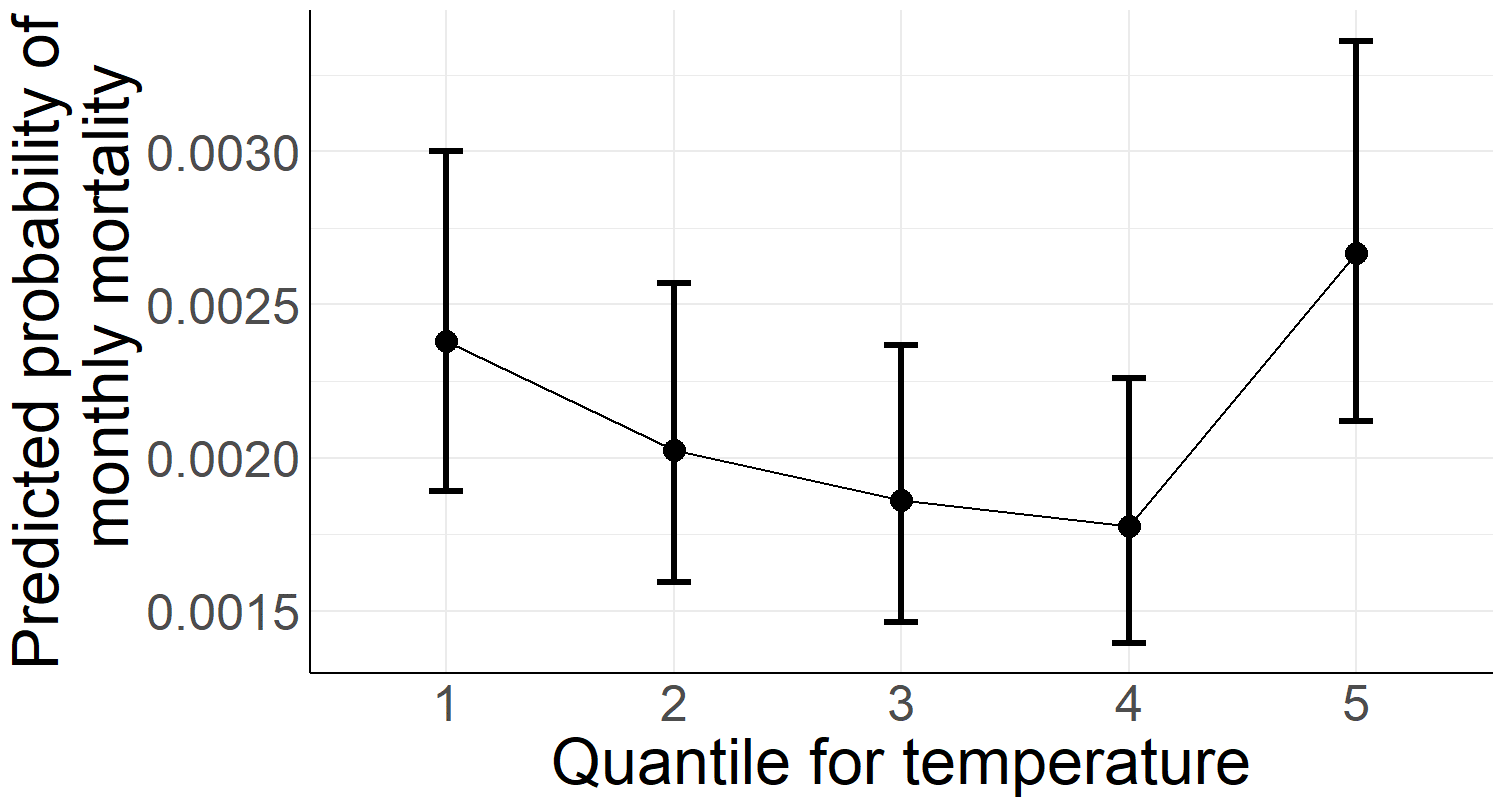 | f)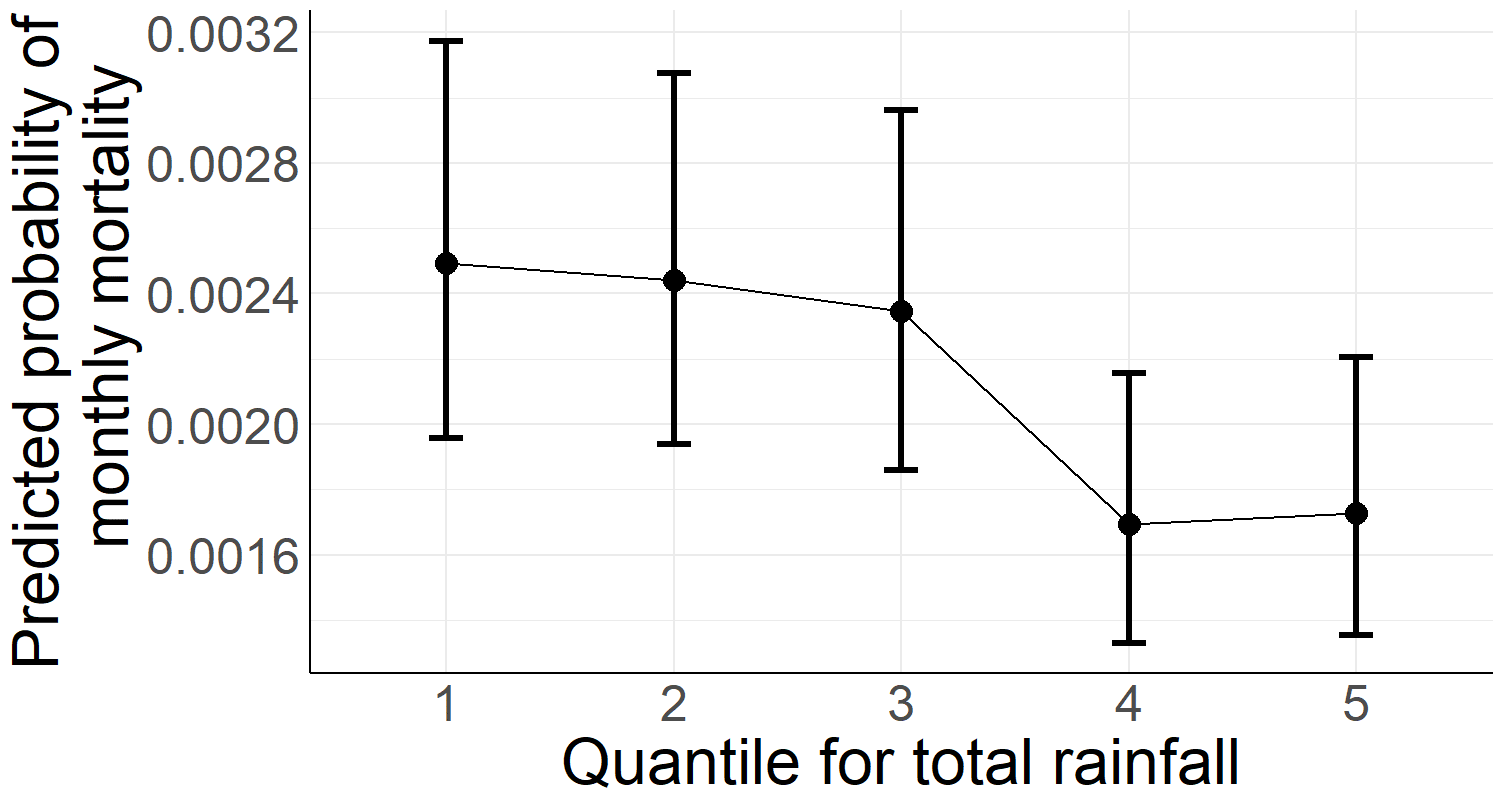 |
| g)  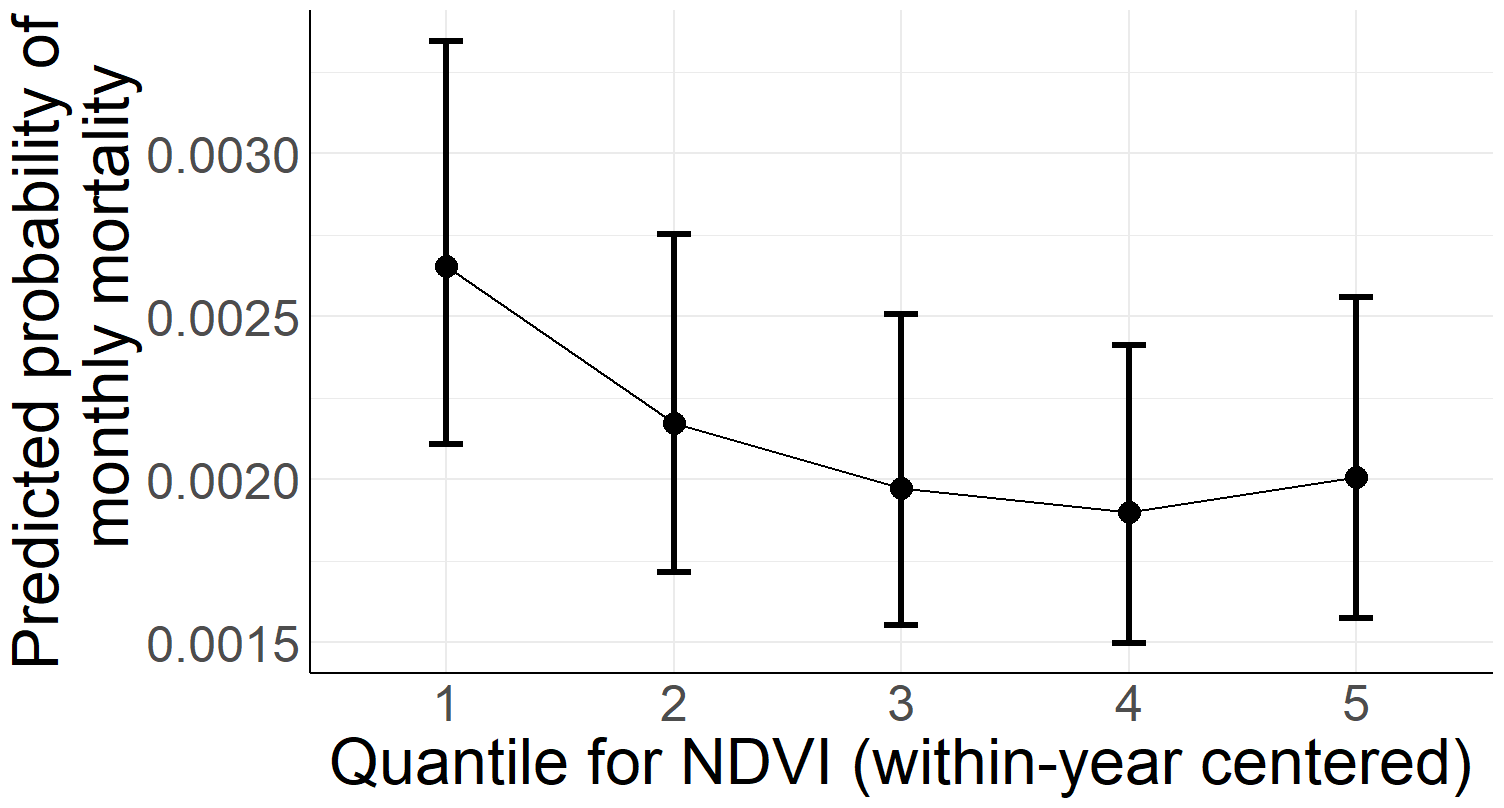 | h)  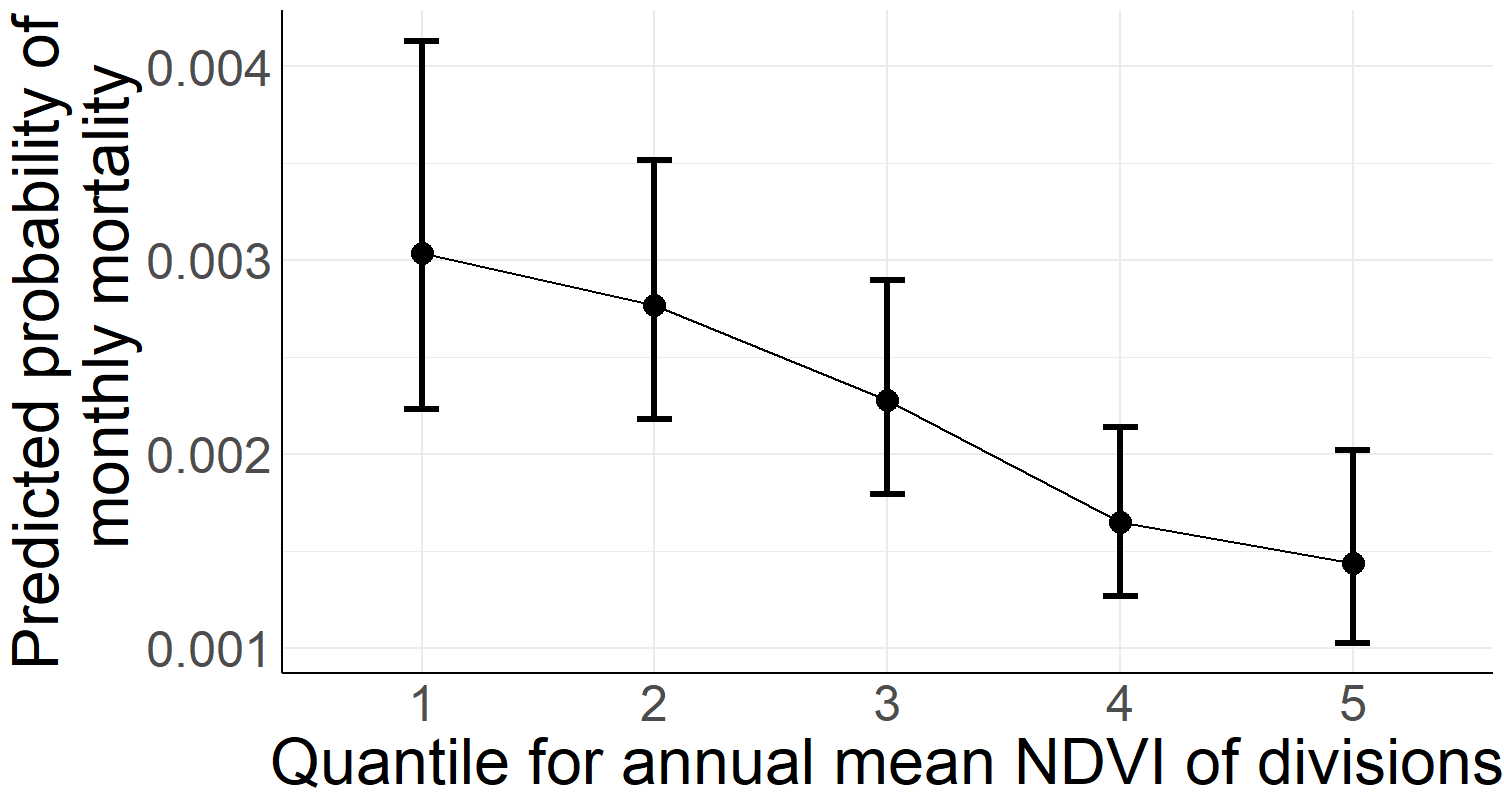 |

Fig. S4. Inspecting non-linearity of the effects of a) temperature, b) rainfall, c) NDVI_div_ and d) NDVI_within-year_ on logit odds of monthly mortality. Plots were obtained from models where the variable of interest was added to the baseline model (see main text). Plots and comparison of means using *emmeans* suggest that U-shaped effect of temperature (middle quantile differs from extreme quantiles 1 and 5, P<0.05) whereas other predictors appear to have largely linear effects (mortality declining with an increase in rainfall, NDVI_within-year_ and NDVI_div_). Plots e-h show same effects converted for odds of monthly mortality as the scale of response.

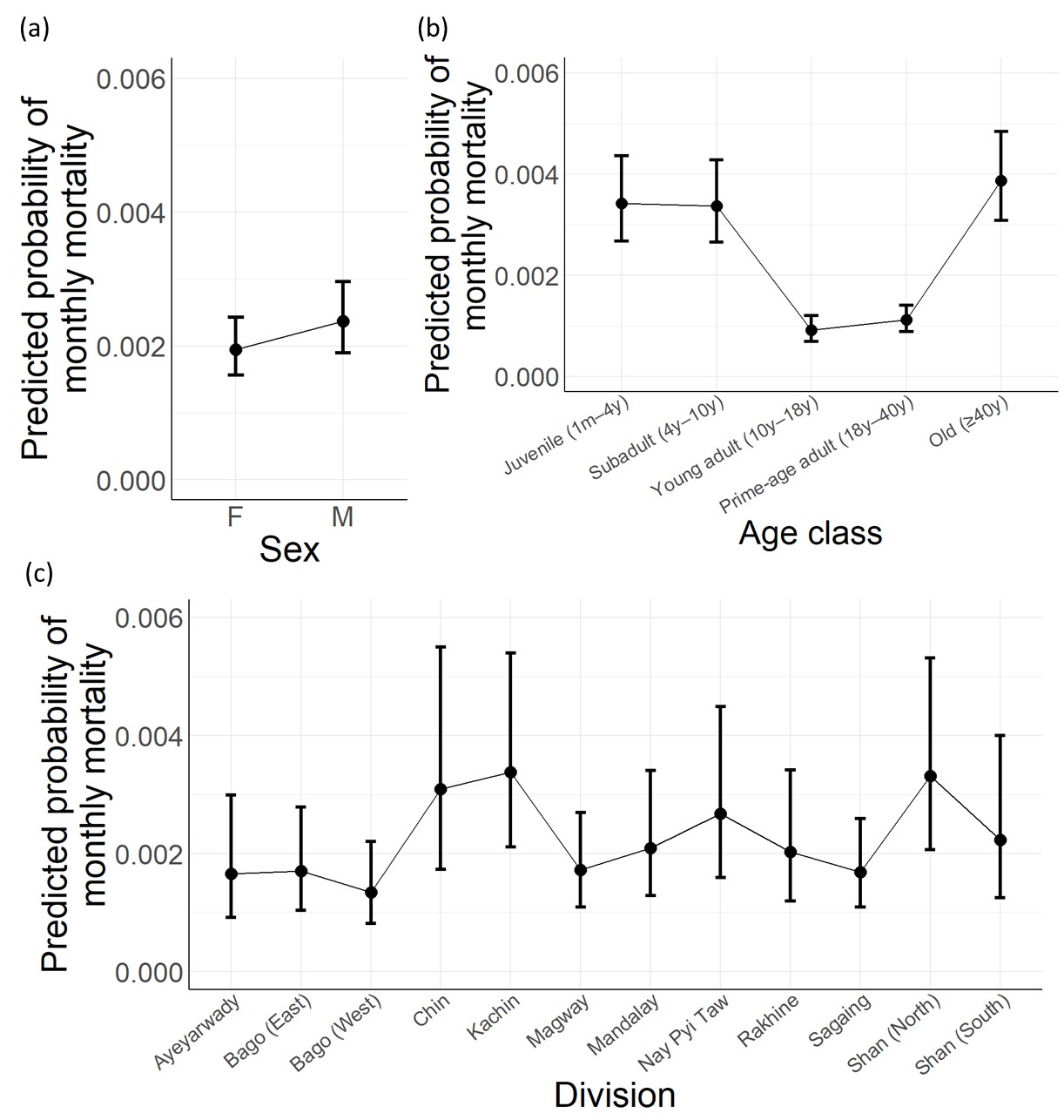

Fig. S5. Predicted probability of monthly mortality obtained from the baseline model reported in the main text. Responses were obtained using *emmeans* (type = response).

**Effects based on competing model ranked 7:** prob.(mortality) ~ b + (t × r) + (t × n_d_) + n_w_

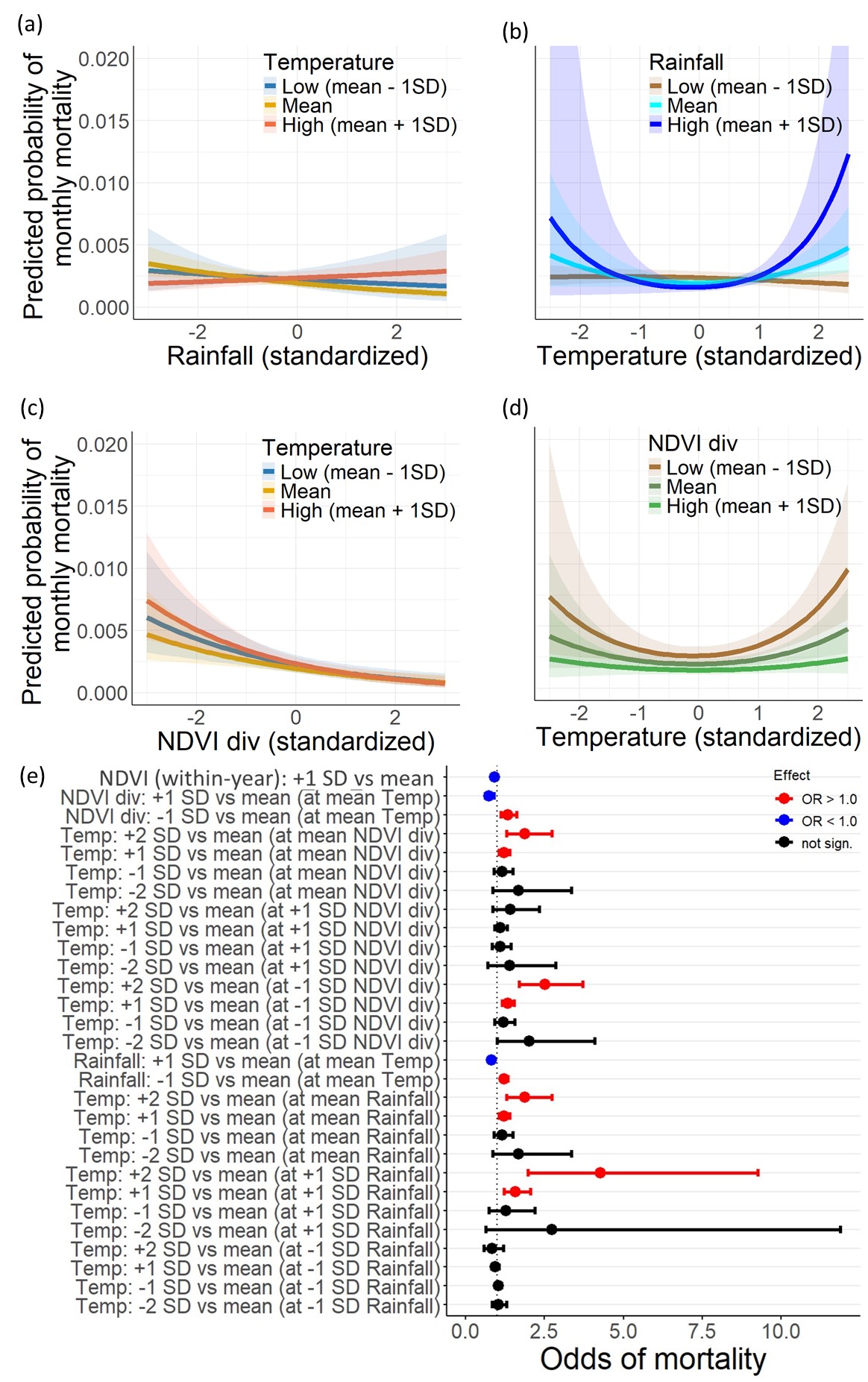

Fig. S6. Effects based on the model containing current-month rainfall and NDVI_within-year_ as one of the predictors (model ranked 7, Table S6). Predicted mortality rates across standardized effects of a) current-month rainfall (response for three discrete temperature values due to interaction effects), b) temperature (response for three discrete rainfall values), c) NDVI_div_ (at three discrete temperature values), d) temperature (response for three discrete NDVI_div_ values), and e) Effect sizes for environmental predictors shown as changes in the odds of mortality. Response curves were obtained using *emmeans* and are averaged across age, sex and divisions.

| **a)**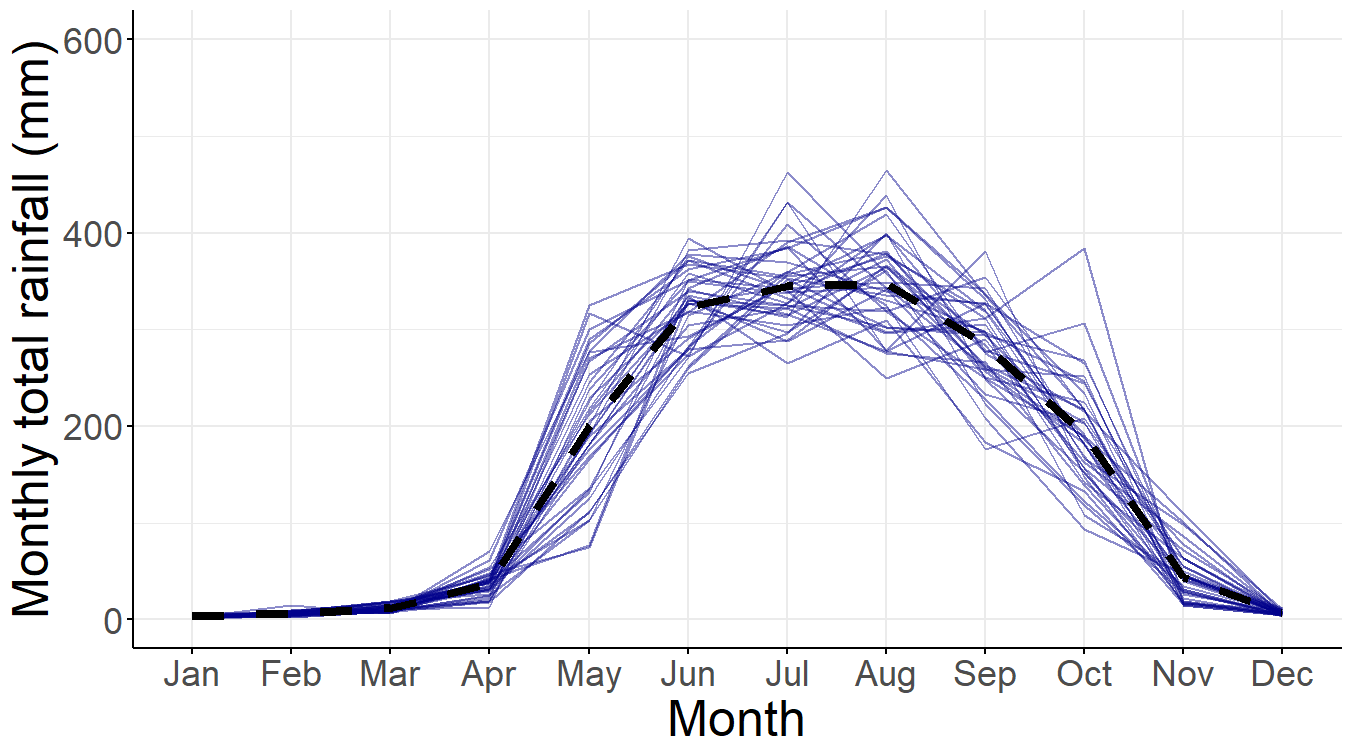 | **b)**  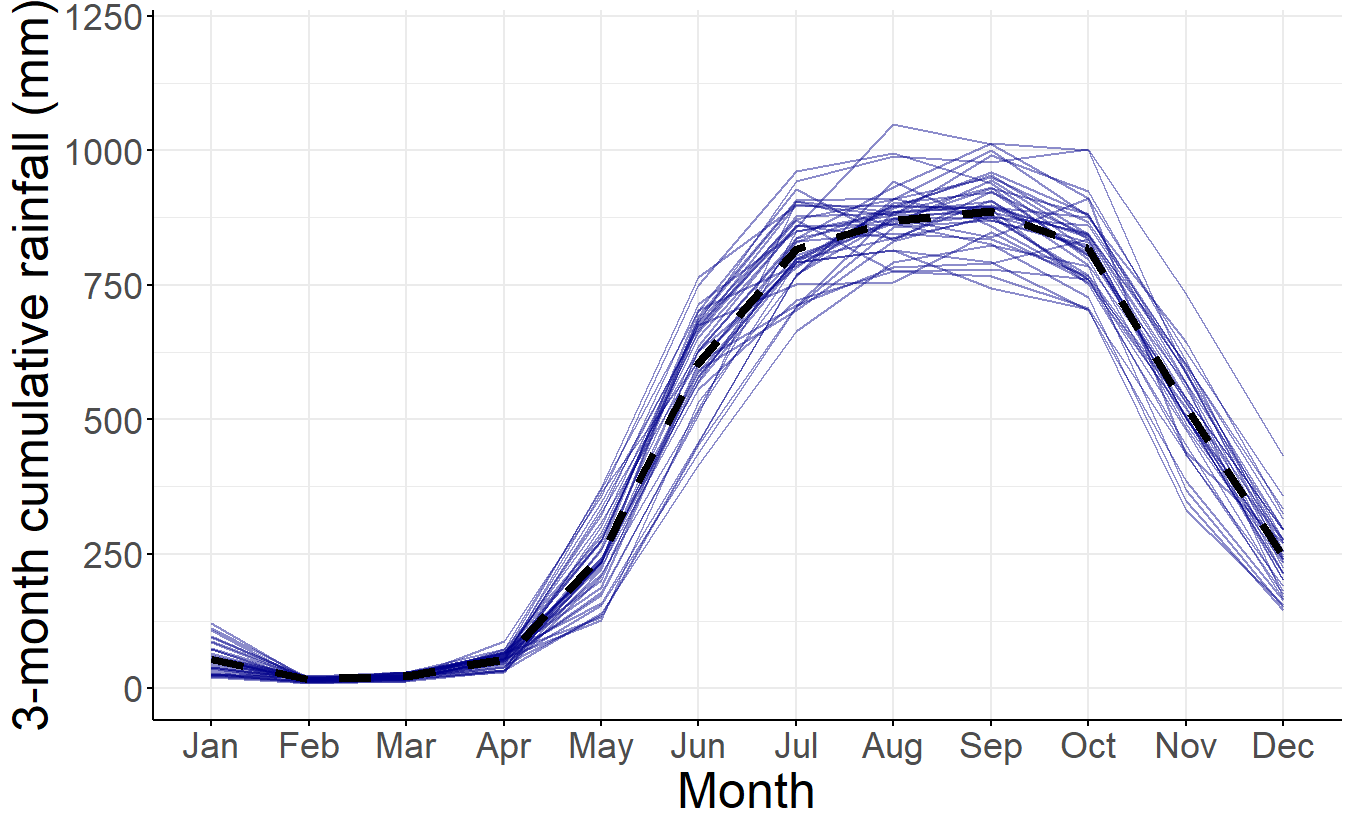 |
| --- | --- |
| **c)**  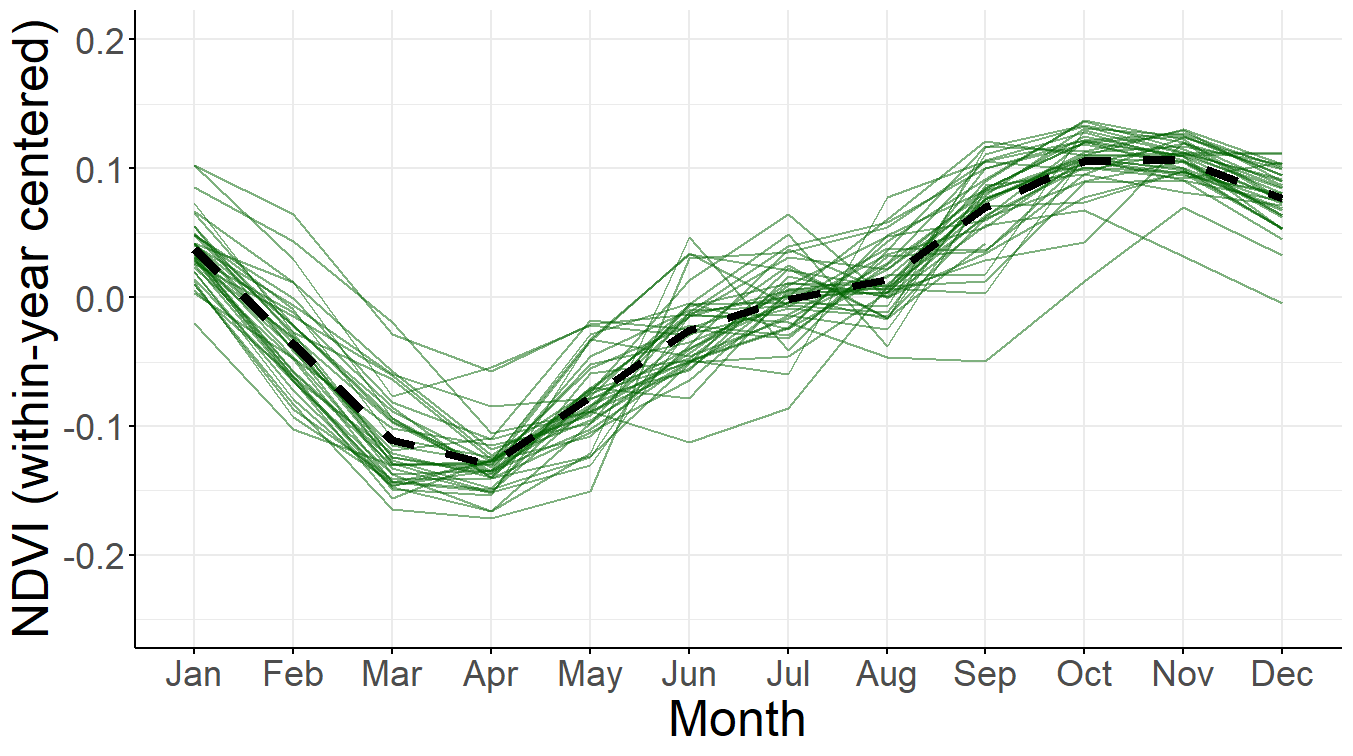 | **d)**  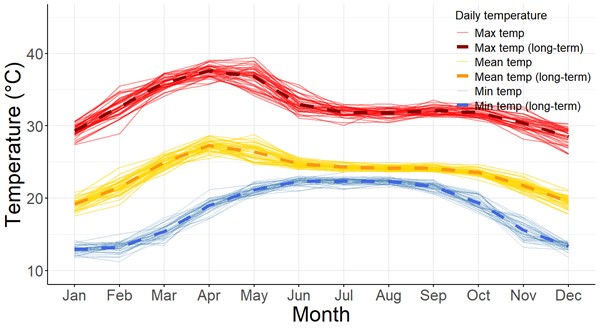 |

Fig. S7. Seasonality plots for a) current-month rainfall b) 3-month cumulative rainfall c) NDVI (within-year centered), and d) ambient temperature (showing monthly averages of mean, minimum and maximum daily temperature). Thin coloured line represents a year and their values are averaged across all 12 divisions considered in this study, and the thick dashed line represents long-term mean across all years.

Tables

Table S1. AIC-based performance of the models with window effects of different variables on mortality. The time window chosen for analyses described in the main text is highlighted for each variable. k = no. of parameters in the model, *k is larger by one for models with temperature because its effects were modelled using spline terms two degrees of freedom.

|  | Current-month measurement | Rolling statistic over 2 months | Rolling statistic over 3 months | Rolling statistic over 4 months |
| --- | --- | --- | --- | --- |
| b + Sum of total rainfall (k=22*) | 12191.29 | 12173.03 | **12164.28** | 12164.26 |
| b + Average of temperature (k=23*) | **12192.48** | 12199.41 | 12207.32 | 12202.87 |
| b + Average of NDVI_within-year_ (k=23*) | **12199.33** | 12205.55 | 12214.28 | 12218.21 |

Table S2. AIC-based comparison of all candidate models of monthly mortality. The baseline model (b) includes the fixed effects of age, sex and division categories, and random effects of year and temporal autocorrelation across months. r = rainfall, r_3_ = total rainfall over 3 months, t = temperature (natural splines, *df*=2), n_d_ = annual mean NDVI of the division, n_w_= within-year centered NDVI (i.e., monthly deviation from n_d_). Top models with ΔAIC ≤ 6.0 are highlighted.

| **Model** | **k** | **AIC** | **ΔAIC** | **AIC_w_** | **Model rank** |
| --- | --- | --- | --- | --- | --- |
| b | 21 | 12216.41 | 69.503 | 0.000 | 26 |
| b + r + t | 24 | 12178.11 | 31.202 | 0.000 | 21 |
| b + r_3_ + t | 24 | 12161.48 | 14.571 | 0.000 | 13 |
| b + (r × t) | 26 | 12172.02 | 25.120 | 0.000 | 19 |
| b + (r_3_ × t) | 26 | 12164.35 | 17.441 | 0.000 | 16 |
| b + t + n_d_ | 24 | 12180.91 | 34.001 | 0.000 | 23 |
| b + (t × n_d_) | 26 | 12178.19 | 31.282 | 0.000 | 22 |
| b + t + n_w_ | 24 | 12183.82 | 36.918 | 0.000 | 24 |
| b + (t × n_w_) | 26 | 12185.78 | 38.871 | 0.000 | 25 |
| b + t + n_d_ + n_w_ | 25 | 12172.01 | 25.103 | 0.000 | 18 |
| b + r + t + n_d_ | 25 | 12166.15 | 19.247 | 0.000 | 17 |
| b + r_3_ + t + n_d_ | **25** | **12149.24** | **2.334** | **0.157** | **2** |
| b + r + (t × n_d_) | 27 | 12164.25 | 17.349 | 0.000 | 15 |
| b + r_3_ + (t × n_d_) | **27** | **12146.9** | **0.000** | **0.505** | **1** |
| b + (r × t) + n_d_ | 27 | 12160.02 | 13.115 | 0.001 | 12 |
| b + (r_3_ × t) + n_d_ | **27** | **12152.09** | **5.186** | **0.038** | **6** |
| b + (r × t) + n_d_ + n_w_ | 28 | 12155.51 | 8.606 | 0.007 | 8 |
| b + (r_3_ × t) + n_d_ + n_w_ | 29 | 12358.54 | 211.634 | 0.000 | 27 |
| b + r + t + n_w_ | 25 | 12174.63 | 27.722 | 0.000 | 20 |
| b + r_3_ + t + n_w_ | 26 | 12368.3 | 221.400 | 0.000 | 28 |
| b + r + t + n_d_ + n_w_ | 26 | 12162.28 | 15.375 | 0.000 | 14 |
| b + r_3_ + t + n_d_ + n_w_ | **26** | **12150.95** | **4.046** | **0.067** | **4** |
| b + (r × t) + (t × n_d_) | 29 | 12157.63 | 10.720 | 0.002 | 10 |
| b + (r_3_ × t) + (t × n_d_) | **29** | **12149.68** | **2.772** | **0.126** | **3** |
| b + (r × t) + (t × n_d_) + n_w_ | **30** | **12152.68** | **5.777** | **0.028** | **7** |
| b + (r_3_ × t) + (t × n_d_) + n_w_ | **30** | **12151.12** | **4.212** | **0.061** | **5** |
| b + (r × t) + n_d_ + (t × n_w_) | 30 | 12157.94 | 11.035 | 0.002 | 11 |
| b + (r_3_ × t) + n_d_ + (t × n_w_) | 30 | 12156.48 | 9.575 | 0.004 | 9 |

**Table S3.** AIC-based performance of the variants of the top seven models of monthly mortality, where different shapes of temperature effects were examined by changing the degrees of freedom of natural splines (df=1–4) or by exploring the quadratic effects of temperature which allow only symmetric U-shaped effects as compared to the flexible shape in the model with two degrees of freedom of natural splines. The asymmetric U shape of temperature effects appears to have marginally better support than the symmetric U-shaped effects (quadratic effects). While the more complex curvature of models with four degrees of freedom fitted well in models ranked 3 and 6, note that these complex curves have two additional parameters (two additional degrees of freedom) but still had AIC within six points in most cases, and thus were not substantially better than the model with df=2. k = no. of parameters in the model.

|  | Natural splines (df=1, i.e., linear effects) | Natural splines (df=2, i.e., asymmetric U shape) | Quadratic effects (i.e., symmetric U shape) | Natural splines (df=3) | Natural splines (df=4) |
| --- | --- | --- | --- | --- | --- |
| Top-ranked model  b + r_3_ + (t × n_d_) | 12153.76 | **12146.90** | **12147.41** | **12149.34** | **12152.66** |
| Model ranked 2  b + r_3_ + t + n_d_ | **12151.80** | **12149.24** | **12149.80** | **12150.77** | **12152.29** |
| Model ranked 3  b + (r_3_ × t) + (t × n_d_) | 12155.65 | **12149.68** | 12150.8 | **12148.94** | **12144.57** |
| Model ranked 4  b + r_3_ + t + nd + nw | **12153.13** | **12150.95** | **12151.50** | **12152.55** | **12154.07** |
| Model ranked 5  b + (r_3_ × t) + (t × n_d_) + n_w_ | 12156.90 | **12151.12** | **12151.63** | **12150.03** | **12146.17** |
| Model ranked 6  b + (r_3_ × t) + n_d_ | 12153.69 | 12152.09 | 12152.58 | 12150.83 | **12144.12** |
| Model ranked 7  b + (r × t) + (t × n_d_) + n_w_ | 12161.29 | **12152.68** | **12152.64** | **12152.71** | **12153.67** |

**Table S4**. Effect sizes expressed as odds of mortality risk for different fixed effects in **a)** **model ranked 4** and its subset **b)** **model ranked 2** that did not contain NDVI_within-year_ (highlighted in the end). In model 4, NDVI_within-year_ neither had a significant effect nor its presence substantially alter the effects of temperature obtained from model 2 that does not contain NDVI_within-year_, thus largely inconsistent with a prediction that NDVI_within-year_ influences the effects of temperature extremes. NDVI_within-year_ only marginally influenced the estimated effects of heat (+1 SD vs mean). The odds of mortality are with respect to mortality at the reference categories or at mean. Other estimated effects also remain largely stable across the models.

| **a) Model 4**:  mortality ~ b + r_3_ + t + n_d_ + n_w_ | | | | | **b) Model 2**:  mortality ~ b + r_3_ + t + n_d_ | | | |
| --- | --- | --- | --- | --- | --- | --- | --- | --- |
| Term | Odds of mortality | 95% CI (low) | 95% CI (high) | *P* | Odds of mortality | 95% CI (low) | 95% CI (high) | *P* |
| Age class (ref: juvenile) | 1.0 | - | - | - | 1.0 | - | - | - |
| Age: subadult | 0.988 | 0.832 | 1.172 | 0.887 | 0.988 | 0.832 | 1.172 | 0.886 |
| **Age: young adult** | **0.267** | **0.214** | **0.333** | **0.000** | **0.267** | **0.214** | **0.333** | **0.000** |
| **Age: prime-aged adult** | **0.327** | **0.279** | **0.385** | **0.000** | **0.327** | **0.279** | **0.385** | **0.000** |
| Age: old adult | 1.137 | 0.969 | 1.334 | 0.114 | 1.137 | 0.969 | 1.334 | 0.115 |
| Sex (ref: female) | 1.0 | - | - | - | 1.0 | - | - | - |
| **Sex: Male** | **1.213** | **1.094** | **1.344** | **0.000** | **1.213** | **1.094** | **1.344** | **0.000** |
| Division (Ref: Ayeyarwaddy) | 1.0 | - | - | - | 1.0 | - | - | - |
| Division: Bago East | 1.333 | 0.632 | 2.813 | 0.450 | 1.330 | 0.630 | 2.808 | 0.454 |
| DivBago West | 0.811 | 0.390 | 1.689 | 0.576 | 0.811 | 0.389 | 1.689 | 0.575 |
| **DivChin** | **2.946** | **1.282** | **6.772** | **0.011** | **3.011** | **1.314** | **6.900** | **0.009** |
| **DivKachin** | **4.042** | **1.832** | **8.917** | **0.001** | **4.097** | **1.859** | **9.029** | **0.000** |
| DivMagway | 1.060 | 0.526 | 2.140 | 0.870 | 1.067 | 0.529 | 2.154 | 0.855 |
| DivMandalay | 1.174 | 0.568 | 2.425 | 0.666 | 1.180 | 0.571 | 2.439 | 0.655 |
| DivNay Pyi Taw | 1.938 | 0.908 | 4.134 | 0.087 | 1.940 | 0.909 | 4.140 | 0.087 |
| DivRakhine | 2.120 | 0.960 | 4.684 | 0.063 | 2.126 | 0.962 | 4.698 | 0.062 |
| DivSagaing | 1.416 | 0.691 | 2.899 | 0.342 | 1.424 | 0.695 | 2.917 | 0.334 |
| **DivShan North** | **2.986** | **1.405** | **6.347** | **0.004** | **3.019** | **1.421** | **6.413** | **0.004** |
| DivShan South | 1.501 | 0.672 | 3.356 | 0.322 | 1.524 | 0.683 | 3.402 | 0.303 |
| Continuous predictor (Ref: mean) | 1.0 | - | - | - |  |  |  |  |
| **3-month rainfall: + 1SD vs mean** | **0.825** | **0.760** | **0.894** | **0.000** | **0.815** | **0.760** | **0.874** | **0.000** |
| **NDVI_div_: +1SD vs mean** | **0.696** | **0.588** | **0.825** | **0.000** | **0.698** | **0.589** | **0.827** | **0.000** |
| ***Temp: +2 SD vs mean** | **1.306** | **1.006** | **1.697** | **0.045** | **1.350** | **1.071** | **1.703** | **0.011** |
| *Temp: +1 SD vs mean | 1.100 | 0.994 | 1.217 | 0.065 | **1.117** | **1.026** | **1.216** | **0.011** |
| *Temp: -1 SD vs mean | 1.003 | 0.936 | 1.076 | 0.927 | 0.992 | 0.938 | 1.049 | 0.771 |
| *Temp: -2 SD vs mean | 1.084 | 0.912 | 1.287 | 0.359 | 1.062 | 0.909 | 1.241 | 0.449 |
| NDVI_within-year_: +1 SD vs mean | 0.980 | 0.909 | 1.056 | 0.590 | - | - | - | - |

Marginal effects for continuous predictors were obtained by generating predicted log-odds using *emmeans*, keeping other effects constant, and then exponentiating to obtain a marginal odds ratio (contrasts obtained using *revpairwise* method and *response* type). *As temperature involves spline terms whose effects cannot be directly inferred from summary table, its predicted marginal effects for up to 2SD difference from mean temperature are shown.

**Table S5.** Effect sizes expressed as odds of mortality risk for different fixed effects in **a) model ranked 3** that did not contain NDVI_within-year_ and **b) model ranked 5** containing this predictor (row highlighted in the end). NDVI_within-year_ neither had a significant effect nor altered the temperature effects obtained from model 3 (without NDVI_within-year_), thus contradicting a prediction that it influences temperature effects. Odds of mortality are with respect to mortality at reference categories or mean.

| **a) Model 3** (AIC = 12149.68):  mortality ~ b + (t × r_3_) + (t × n_d_) | | | | | **b) Model 5** (12151.12):  mortality ~ b + (t × r_3_) + (t × n_d_) + n_w_ | | | |
| --- | --- | --- | --- | --- | --- | --- | --- | --- |
| Term | Odds of mortality | 95% CI (low) | 95% CI (high) | *P* | Odds of mortality | 95% CI (low) | 95% CI (high) | *P* |
| Age class (ref: juvenile) | 1.0 | - | - | - | 1.0 | - | - | - |
| Age: subadult | 0.987 | 0.832 | 1.171 | 0.880 | 0.987 | 0.832 | 1.171 | 0.881 |
| **Age: young adult** | **0.267** | **0.214** | **0.333** | **0.000** | **0.267** | **0.214** | **0.333** | **0.000** |
| **Age: prime-aged adult** | **0.327** | **0.278** | **0.384** | **0.000** | **0.327** | **0.278** | **0.384** | **0.000** |
| Age: old adult | 1.136 | 0.968 | 1.332 | 0.119 | 1.136 | 0.968 | 1.333 | 0.119 |
| Sex (ref: female) | 1.0 | - | - | - | 1.0 | - | - | - |
| **Sex: Male** | **1.213** | **1.094** | **1.344** | **0.000** | **1.213** | **1.094** | **1.344** | **0.000** |
| Division (Ref: Ayeyarwaddy) | 1.0 | - | - | - | 1.0 | - | - | - |
| Division: Bago East | 1.370 | 0.649 | 2.892 | 0.409 | 1.375 | 0.652 | 2.903 | 0.403 |
| DivBago West | 0.819 | 0.394 | 1.706 | 0.594 | 0.821 | 0.394 | 1.707 | 0.597 |
| **DivChin** | **2.969** | **1.286** | **6.854** | **0.011** | **2.860** | **1.232** | **6.636** | **0.014** |
| **DivKachin** | **4.095** | **1.854** | **9.042** | **0.000** | **4.015** | **1.816** | **8.876** | **0.001** |
| DivMagway | 1.081 | 0.536 | 2.182 | 0.827 | 1.071 | 0.531 | 2.161 | 0.848 |
| DivMandalay | 1.187 | 0.575 | 2.452 | 0.643 | 1.177 | 0.570 | 2.432 | 0.659 |
| DivNay Pyi Taw | 1.974 | 0.924 | 4.215 | 0.079 | 1.971 | 0.923 | 4.207 | 0.080 |
| DivRakhine | 2.116 | 0.956 | 4.681 | 0.064 | 2.110 | 0.954 | 4.666 | 0.065 |
| DivSagaing | 1.421 | 0.694 | 2.906 | 0.336 | 1.409 | 0.689 | 2.881 | 0.348 |
| **DivShan North** | **3.009** | **1.416** | **6.397** | **0.004** | **2.961** | **1.392** | **6.300** | **0.005** |
| DivShan South | 1.505 | 0.673 | 3.369 | 0.320 | 1.470 | 0.655 | 3.295 | 0.350 |
| Continuous predictor (Ref: mean) | 1.0 | - | - | - | 1.0 | - | - | - |
| NDVI_div_: +1SD (at mean temp) | **0.747** | **0.625** | **0.895** | **0.002** | **0.746** | **0.623** | **0.893** | **0.001** |
| NDVI_div_: - 1SD (at mean temp) | **1.338** | **1.118** | **1.601** | **0.002** | **1.341** | **1.120** | **1.604** | **0.001** |
| *Temp: +2 SD (at mean NDVI_div_) | 1.537 | 0.956 | 2.471 | 0.076 | 1.475 | 0.907 | 2.399 | 0.117 |
| *Temp: +1 SD (at mean NDVI_div_) | 1.159 | 0.977 | 1.374 | 0.090 | 1.135 | 0.949 | 1.357 | 0.166 |
| *Temp: -1 SD (at mean NDVI_div_) | 1.026 | 0.922 | 1.142 | 0.635 | 1.050 | 0.929 | 1.186 | 0.436 |
| *Temp: -2 SD (at mean NDVI_div_) | 1.200 | 0.877 | 1.641 | 0.255 | 1.256 | 0.899 | 1.756 | 0.182 |
| *Temp: +2 SD (at +1 SD NDVI_div_) | 1.161 | 0.665 | 2.027 | 0.599 | 1.111 | 0.629 | 1.964 | 0.716 |
| *Temp: +1 SD (at +1 SD NDVI_div_) | 1.059 | 0.866 | 1.296 | 0.577 | 1.036 | 0.840 | 1.278 | 0.742 |
| *Temp: -1 SD (at +1 SD NDVI_div_) | 0.987 | 0.867 | 1.124 | 0.844 | 1.010 | 0.875 | 1.164 | 0.896 |
| *Temp: -2 SD (at +1 SD NDVI_div_) | 1.007 | 0.695 | 1.459 | 0.970 | 1.054 | 0.715 | 1.555 | 0.791 |
| ***Temp: +2 SD (at -1 SD NDVI_div_)** | **2.034** | **1.217** | **3.399** | **0.007** | **1.958** | **1.161** | **3.303** | **0.012** |
| ***Temp: +1 SD (at -1 SD NDVI_div_)** | **1.268** | **1.055** | **1.525** | **0.012** | **1.243** | **1.027** | **1.505** | **0.026** |
| *Temp: -1 SD (at -1 SD NDVI_div_) | 1.067 | 0.948 | 1.201 | 0.281 | 1.091 | 0.957 | 1.245 | 0.194 |
| ***Temp: -2 SD (at -1 SD NDVI_div_)** | **1.429** | **1.012** | **2.017** | **0.042** | **1.497** | **1.039** | **2.156** | **0.030** |
| **3-month rainfall: +1 SD (at mean temp)** | **0.796** | **0.729** | **0.868** | **0.000** | **0.809** | **0.735** | **0.890** | **0.000** |
| **3-month rainfall: -1 SD (at mean Temp)** | **1.257** | **1.152** | **1.371** | **0.000** | **1.237** | **1.124** | **1.361** | **0.000** |
| *Temp: +2 SD (at mean 3m rainfall) | 1.537 | 0.956 | 2.471 | 0.076 | 1.475 | 0.907 | 2.399 | 0.117 |
| *Temp: +1 SD (at mean 3m rainfall) | 1.159 | 0.977 | 1.374 | 0.090 | 1.135 | 0.949 | 1.357 | 0.166 |
| *Temp: -1 SD (at mean 3m rainfall) | 1.026 | 0.922 | 1.142 | 0.635 | 1.050 | 0.929 | 1.186 | 0.436 |
| *Temp: -2 SD (at mean 3m rainfall) | 1.200 | 0.877 | 1.641 | 0.255 | 1.256 | 0.899 | 1.756 | 0.182 |
| *Temp: +2 SD (at +1 SD 3m rainfall) | 2.048 | 0.758 | 5.533 | 0.158 | 1.982 | 0.732 | 5.370 | 0.178 |
| *Temp: +1 SD (at +1 SD 3m rainfall) | 1.270 | 0.891 | 1.812 | 0.187 | 1.245 | 0.869 | 1.782 | 0.232 |
| *Temp: -1 SD (at +1 SD 3m rainfall) | 1.070 | 0.848 | 1.350 | 0.568 | 1.104 | 0.863 | 1.411 | 0.431 |
| *Temp: -2 SD (at +1 SD 3m rainfall) | 1.441 | 0.730 | 2.846 | 0.292 | 1.546 | 0.765 | 3.122 | 0.225 |
| *Temp: +2 SD (at -1 SD 3m rainfall) | 1.153 | 0.856 | 1.554 | 0.349 | 1.098 | 0.793 | 1.519 | 0.573 |
| *Temp: +1 SD (at -1 SD 3m rainfall) | 1.057 | 0.947 | 1.180 | 0.320 | 1.035 | 0.915 | 1.171 | 0.588 |
| *Temp: -1 SD (at -1 SD 3m rainfall) | 0.984 | 0.917 | 1.057 | 0.668 | 0.998 | 0.921 | 1.081 | 0.965 |
| *Temp: -2 SD (at -1 SD 3m rainfall) | 0.999 | 0.820 | 1.216 | 0.990 | 1.021 | 0.832 | 1.253 | 0.843 |
| NDVI_within-year_: +1 SD | - | - | - | - | 0.971 | 0.900 | 1.048 | 0.452 |

Marginal effects for continuous predictors were obtained by generating predicted log-odds using *emmeans*, keeping other effects constant, and then exponentiating to obtain a marginal odds ratio (contrasts obtained using *revpairwise* method and *response* type). *As temperature involves interactions and spline terms, its predicted marginal effects (for up to 2SD difference from mean temperature) are shown at three discrete values of NDVI_div_ and 3-month rainfall.

**Table S6.** Effect sizes expressed as odds of mortality risk for different fixed effects in model ranked 6 (prob. mortality ~ b + (t × r_3_) + n_d_). The odds of mortality are with respect to mortality at reference categories (categorical predictors) or at the mean value (continuous predictors).

| Term | Odds of mortality | 95% CI (low) | 95% CI (high) | p_value |
| --- | --- | --- | --- | --- |
| Age class (ref: juvenile) | 1.0 | - | - | - |
| Age: subadult | 0.986 | 0.831 | 1.170 | 0.869 |
| **Age: young adult** | **0.267** | **0.214** | **0.333** | **0.000** |
| **Age: prime-aged adult** | **0.328** | **0.279** | **0.385** | **0.000** |
| Age: old adult | 1.136 | 0.968 | 1.333 | 0.119 |
| Sex (ref: female) | 1.0 | - | - | - |
| **Sex: Male** | **1.213** | **1.094** | **1.345** | **0.000** |
| Division (Ref: Ayeyarwaddy) | 1.0 | - | - | - |
| Division: Bago East | 1.443 | 0.686 | 3.036 | 0.334 |
| DivBago West | 0.874 | 0.421 | 1.813 | 0.718 |
| **DivChin** | **3.355** | **1.432** | **7.857** | **0.005** |
| **DivKachin** | **4.510** | **2.046** | **9.943** | **0.000** |
| DivMagway | 1.165 | 0.580 | 2.339 | 0.669 |
| DivMandalay | 1.294 | 0.629 | 2.661 | 0.484 |
| **DivNay Pyi Taw** | **2.166** | **1.018** | **4.606** | **0.045** |
| **DivRakhine** | **2.284** | **1.036** | **5.036** | **0.041** |
| DivSagaing | 1.493 | 0.732 | 3.045 | 0.270 |
| **DivShan North** | **3.357** | **1.584** | **7.112** | **0.002** |
| DivShan South | 1.701 | 0.759 | 3.814 | 0.197 |
| Continuous predictor (Ref: mean) | 1.0 | - | - | - |
| **NDVI_div_: +1SD vs mean** | **0.477** | **0.321** | **0.707** | **0.000** |
| **3-month rainfall: +1 SD (at mean temp)** | **0.799** | **0.733** | **0.872** | **0.000** |
| **3-month rainfall: -1 SD (at mean Temp)** | **1.251** | **1.146** | **1.365** | **0.000** |
| ***Temp: +2 SD (at mean 3m rainfall)** | **1.604** | **1.000** | **2.572** | **0.050** |
| *Temp: +1 SD (at mean 3m rainfall) | 1.175 | 0.992 | 1.392 | 0.061 |
| *Temp: -1 SD (at mean 3m rainfall) | 1.031 | 0.927 | 1.146 | 0.579 |
| *Temp: -2 SD (at mean 3m rainfall) | 1.227 | 0.898 | 1.675 | 0.199 |
| *Temp: +2 SD (at +1 SD 3m rainfall) | 2.010 | 0.744 | 5.432 | 0.169 |
| *Temp: +1 SD (at +1 SD 3m rainfall) | 1.255 | 0.880 | 1.790 | 0.210 |
| *Temp: -1 SD (at +1 SD 3m rainfall) | 1.090 | 0.866 | 1.373 | 0.462 |
| *Temp: -2 SD (at +1 SD 3m rainfall) | 1.505 | 0.765 | 2.959 | 0.236 |
| *Temp: +2 SD (at -1 SD 3m rainfall) | 1.280 | 0.973 | 1.683 | 0.078 |
| *Temp: +1 SD (at -1 SD 3m rainfall) | 1.101 | 0.996 | 1.217 | 0.059 |
| *Temp: -1 SD (at -1 SD 3m rainfall) | 0.974 | 0.908 | 1.045 | 0.464 |
| *Temp: -2 SD (at -1 SD 3m rainfall) | 1.000 | 0.822 | 1.216 | 1.000 |

Marginal effects for continuous predictors were obtained by generating predicted log-odds using *emmeans*, keeping other effects constant, and then exponentiating to obtain a marginal odds ratio (contrasts obtained using *revpairwise* method and *response* type). *As temperature involves interactions and spline terms, its predicted marginal effects (for up to 2SD difference from mean temperature) are shown at three discrete values of 3-month rainfall.

**Table S7**. Effect sizes expressed as odds of mortality risk for different fixed effects in **a) model ranked 7** and its subset **b) model ranked 10** that did not contain NDVI_within-year_. In model 7, NDVI_within-year_ had a significant effect (single-term deletion test, *P*=0.008) but its presence only marginally reduced the estimated effects of heat but not of cold (and only at high NDVI_div_) compared to model 10 that does not contain NDVI_within-year_, thus providing mixed and weak support to the prediction that NDVI_within-year_ influences the effects of temperature. Odds of mortality are with respect to mortality at reference categories or at mean.

| **a) Model 7** (AIC= 12152.68):  mortality ~ b + (t × r) + (t × n_d_) + nw | | | | | **b) Model 10** (AIC= 12157.63):  mortality ~ b + (t × r) + (t × n_d_) | | | |
| --- | --- | --- | --- | --- | --- | --- | --- | --- |
| Term | Odds of mortality | 95% CI (low) | 95% CI (high) | p_value | Odds of mortality | 95% CI (low) | 95% CI (high) | *P* |
| Age class (ref: juvenile) | 1.0 | - | - | - | 1.0 | - | - | - |
| Age: subadult | 0.986 | 0.831 | 1.171 | 0.875 | 0.986 | 0.831 | 1.170 | 0.869 |
| **Age: young adult** | **0.267** | **0.214** | **0.333** | **0.000** | **0.267** | **0.214** | **0.333** | **0.000** |
| **Age: prime-aged adult** | **0.327** | **0.279** | **0.385** | **0.000** | **0.328** | **0.279** | **0.385** | **0.000** |
| Age: old adult | 1.138 | 0.970 | 1.335 | 0.113 | 1.137 | 0.969 | 1.334 | 0.116 |
| Sex (ref: female) | 1.0 | - | - | - | 1.0 | - | - | - |
| **Sex: Male** | **1.213** | **1.094** | **1.344** | **0.000** | **1.213** | **1.094** | **1.345** | **0.000** |
| Division (Ref: Ayeyarwaddy) | 1.0 | - | - | - | 1.0 | - | - | - |
| Division: Bago East | 1.464 | 0.696 | 3.078 | 0.315 | 1.443 | 0.686 | 3.036 | 0.334 |
| DivBago West | 0.875 | 0.422 | 1.814 | 0.720 | 0.874 | 0.421 | 1.813 | 0.718 |
| **DivChin** | **2.986** | **1.270** | **7.020** | **0.012** | **3.355** | **1.432** | **7.857** | **0.005** |
| **DivKachin** | **4.226** | **1.916** | **9.322** | **0.000** | **4.510** | **2.046** | **9.943** | **0.000** |
| DivMagway | 1.123 | 0.559 | 2.256 | 0.744 | 1.165 | 0.580 | 2.339 | 0.669 |
| DivMandalay | 1.250 | 0.607 | 2.571 | 0.545 | 1.294 | 0.629 | 2.661 | 0.484 |
| **DivNay Pyi Taw** | **2.134** | **1.003** | **4.537** | **0.049** | **2.166** | **1.018** | **4.606** | **0.045** |
| **DivRakhine** | **2.248** | **1.020** | **4.954** | **0.044** | **2.284** | **1.036** | **5.036** | **0.041** |
| DivSagaing | 1.448 | 0.711 | 2.952 | 0.308 | 1.493 | 0.732 | 3.045 | 0.270 |
| **DivShan North** | **3.162** | **1.491** | **6.703** | **0.003** | **3.357** | **1.584** | **7.112** | **0.002** |
| DivShan South | 1.554 | 0.692 | 3.493 | 0.286 | 1.701 | 0.759 | 3.814 | 0.197 |
| Continuous predictor (Ref: mean) | 1.0 | - | - | - | 1.0 | - | - | - |
| **NDVI_div_: +1SD (at mean temp)** | **0.745** | **0.623** | **0.892** | **0.001** | **0.749** | **0.625** | **0.897** | **0.002** |
| **NDVI_div_: - 1SD (at mean temp)** | **1.342** | **1.121** | **1.606** | **0.001** | **1.335** | **1.115** | **1.599** | **0.002** |
| ***Temp: +2 SD (at mean NDVI_div_)** | **1.881** | **1.292** | **2.737** | **0.001** | **2.241** | **1.571** | **3.195** | **0.000** |
| ***Temp: +1 SD (at mean NDVI_div_)** | **1.208** | **1.054** | **1.384** | **0.007** | **1.316** | **1.165** | **1.486** | **0.000** |
| *Temp: -1 SD (at mean NDVI_div_) | 1.147 | 0.887 | 1.485 | 0.296 | 1.059 | 0.822 | 1.364 | 0.658 |
| *Temp: -2 SD (at mean NDVI_div_) | 1.682 | 0.846 | 3.346 | 0.138 | 1.438 | 0.726 | 2.850 | 0.298 |
| *Temp: +2 SD (at +1 SD NDVI_div_) | 1.410 | 0.853 | 2.331 | 0.181 | **1.698** | **1.047** | **2.753** | **0.032** |
| *Temp: +1 SD (at +1 SD NDVI_div_) | 1.101 | 0.917 | 1.322 | 0.300 | **1.204** | **1.015** | **1.428** | **0.033** |
| *Temp: -1 SD (at +1 SD NDVI_div_) | 1.101 | 0.844 | 1.437 | 0.477 | 1.018 | 0.784 | 1.322 | 0.894 |
| *Temp: -2 SD (at +1 SD NDVI_div_) | 1.402 | 0.687 | 2.859 | 0.353 | 1.207 | 0.593 | 2.455 | 0.603 |
| ***Temp: +2 SD (at -1 SD NDVI_div_)** | **2.509** | **1.697** | **3.709** | **0.000** | **2.957** | **2.033** | **4.300** | **0.000** |
| ***Temp: +1 SD (at -1 SD NDVI_div_)** | **1.325** | **1.149** | **1.528** | **0.000** | **1.438** | **1.263** | **1.638** | **0.000** |
| *Temp: -1 SD (at -1 SD NDVI_div_) | 1.196 | 0.917 | 1.558 | 0.186 | 1.102 | 0.849 | 1.429 | 0.466 |
| *Temp: -2 SD (at -1 SD NDVI_div_) | 2.018 | 0.997 | 4.088 | 0.051 | 1.714 | 0.850 | 3.455 | 0.132 |
| **Rainfall: +1 SD (at mean temp)** | **0.822** | **0.748** | **0.903** | **0.000** | **0.798** | **0.727** | **0.877** | **0.000** |
| **Rainfall: -1 SD (at mean Temp)** | **1.217** | **1.107** | **1.337** | **0.000** | **1.252** | **1.140** | **1.376** | **0.000** |
| ***Temp: +2 SD (at mean rainfall)** | **1.881** | **1.292** | **2.737** | **0.001** | **2.241** | **1.571** | **3.195** | **0.000** |
| ***Temp: +1 SD (at mean rainfall)** | **1.208** | **1.054** | **1.384** | **0.007** | **1.316** | **1.165** | **1.486** | **0.000** |
| *Temp: -1 SD (at mean rainfall) | 1.147 | 0.887 | 1.485 | 0.296 | 1.059 | 0.822 | 1.364 | 0.658 |
| *Temp: -2 SD (at mean rainfall) | 1.682 | 0.846 | 3.346 | 0.138 | 1.438 | 0.726 | 2.850 | 0.298 |
| ***Temp: +2 SD (at +1 SD rainfall)** | **4.271** | **1.972** | **9.249** | **0.000** | **4.964** | **2.292** | **10.751** | **0.000** |
| ***Temp: +1 SD (at +1 SD rainfall)** | **1.576** | **1.207** | **2.059** | **0.001** | **1.720** | **1.324** | **2.235** | **0.000** |
| *Temp: -1 SD (at +1 SD rainfall) | 1.273 | 0.738 | 2.194 | 0.385 | 1.131 | 0.658 | 1.945 | 0.656 |
| *Temp: -2 SD (at +1 SD rainfall) | 2.732 | 0.629 | 11.869 | 0.180 | 2.108 | 0.486 | 9.155 | 0.319 |
| *Temp: +2 SD (at -1 SD rainfall) | 0.828 | 0.575 | 1.192 | 0.311 | 1.011 | 0.724 | 1.412 | 0.947 |
| *Temp: +1 SD (at -1 SD rainfall) | 0.926 | 0.806 | 1.062 | 0.272 | 1.007 | 0.890 | 1.139 | 0.918 |
| *Temp: -1 SD (at -1 SD rainfall) | 1.034 | 0.951 | 1.125 | 0.430 | 0.991 | 0.917 | 1.072 | 0.827 |
| *Temp: -2 SD (at -1 SD rainfall) | 1.036 | 0.834 | 1.285 | 0.751 | 0.981 | 0.793 | 1.214 | 0.860 |
| **NDVI_within-year_: +1 SD vs mean** | **0.914** | **0.855** | **0.977** | **0.008** | - | - | - | - |

Marginal effects for continuous predictors were obtained by generating predicted log-odds using *emmeans*, keeping other effects constant, and then exponentiating to obtain a marginal odds ratio (contrasts obtained using *revpairwise* method and *response* type). *As temperature involves interactions and spline terms, its predicted marginal effects (for up to 2SD difference from mean temperature) are shown at three discrete values of NDVI_div_ and rainfall.
